# Evaluating the impact of a sample-matched reference genome on single-cell transcriptomic inferences in *Plasmodium falciparum*

**DOI:** 10.64898/2026.07.30.740912

**Authors:** Talleh Almelli, Sunil Kumar Dogga, Jesse Rop, Seri Kitada, Mara Lawniczak

## Abstract

**Background:** *Plasmodium falciparum* field isolates exhibit genomic variation, including copy number variation and sequence divergence. In contrast, the *P. falciparum* 3D7 reference genome (Pf3D7) was derived from a long-term laboratory-adapted strain and does not fully reflect the genomic variation among field isolates. The extent to which the reference genome influences RNA-seq mapping and expression inference in natural infections remains unclear.

**Results:** We generated both a reference genome and single cell RNA sequencing (scRNAseq) data from a *P. falciparum-*infected carrier in Mali. This new ML52 assembly was annotated using Companion with Pf3D7 as the reference. scRNAseq reads from the natural infection isolate were aligned to both the Pf3D7 genome and the isolate-specific ML52 genome, followed by locus-level alignment inspection. For most conserved genes, expression inference was concordant for both references, while genes showing reference-genome-dependent differences were investigated further. Some discrepancies were attributed to mapping artefacts or to reads aligning to unplaced genomic contigs that reflected divergent haplotypes from the co-infecting strains in the naturally infected carrier. While most multigene family loci showed concordant gene expression across both references, *var* genes exhibited considerable mis-mapping against 3D7 *var* loci as well as 2/3 of *var* reads not mapping at all to 3D7.

**Conclusion:** scRNAseq expression inference in *P. falciparum* is robust for conserved genes and most multigene families when comparing to a matched vs unmatched reference genome, but the extremely polymorphic *var* genes require a matched assembly in order to evaluate expression. These findings highlight the importance of the reference genome for *var* gene studies and should be considered when interpreting transcriptomic analyses in other organisms possessing highly variable antigenic loci.

## Introduction

### Genetic and Structural Diversity in *Plasmodium falciparum*

*Plasmodium falciparum* demonstrates genetic and structural diversity across field isolates from natural infections, including single-nucleotide polymorphisms (SNPs), insertions and deletions, gene copy number variation (CNV), and large-scale structural rearrangements [1–3]. These variations are especially prominent in subtelomeric regions, which are also rich in multigene families [4–6]. 3D7 is a laboratory-adapted strain derived from the NF54 isolate, taken from a Dutch patient in the late 1970s who had never traveled outside the country, making it a likely case of “airport malaria.” [7]. The exact origin of NF54 remains unknown, but it is believed to be from Africa [8]. NF54 was cloned through limiting dilution to generate Pf3D7 and has been kept in continuous lab culture for several decades. This extended cultivation led 3D7 to become a single, genetically homogenous clone with reduced intra-isolate diversity and characteristics suited for laboratory culture [9]. Nevertheless, 3D7 remains the principal reference genome for most genomic investigations of *P. falciparum*, including single-cell RNA sequencing (scRNAseq).

The haploid nuclear genome of *P. falciparum* comprises 14 linear chromosomes totalling approximately 23.3 Mb [10]. It is highly AT-rich (∼80.6% overall, up to ∼90% in intronic and intergenic regions) [4,11]. As of VEuPathDB Build 68 (01 September 2020), the GenBank annotation of *Plasmodium falciparum* 3D7 comprised 5,797 genes, of which 5,387 were protein-coding [12]. Comparative analyses across *Plasmodium* species reveal strong conservation in chromosomal core regions but marked variability in subtelomeric domains [13]. These subtelomeric regions consist of repetitive sequence blocks adjacent to telomeric repeats. Six non-coding telomere-associated repeat elements (TAREs) are positioned proximal to the telomeres [5]. Immediately internal to these repeats lie members of the *var* gene family encoding PfEMP1 [14], interspersed by *rif* and *stevor* genes encoding RIFINs [15] and STEVORs [16], respectively. Additional subtelomeric multigene families include *etramp*, *surfin*, *phist*, *clag*, and *fikk* [17].

Major multigene families *var* (∼60 genes in Pf3D7), *rif* (∼149), *stevor* (∼28), and *phist* (∼89) are predominantly subtelomeric and exhibit extensive copy number and structural polymorphism among isolates [18,19]. These genes encode proteins central to cytoadherence, immune evasion, and host–parasite interactions. Their expression is tightly regulated with mutually exclusive transcription of a single *var* gene per parasite at a given time [20,21].

### Single-cell RNA sequencing in *P. falciparum*

Single-cell transcriptomics has become an important approach for studying transcriptional heterogeneity in *P. falciparum*. ScRNAseq studies revealed distinct expression signatures, developmental trajectories, and stage-specific transcriptional programs within parasite populations [22,23]. More recently, comprehensive single-cell atlases spanning intraerythrocytic asexual and sexual development stages have been established, providing high-resolution frameworks for transcriptional programs in laboratory strains and clinical isolates [24,25].

### Technical Considerations in Mapping and Expression Studies

Mapping and quantification steps in short-read scRNAseq inherently depend on aligning to a chosen reference genome and annotation model, which creates sensitivity to the completeness and accuracy of those resources [26]. Mapping biases can arise when the reference diverges in sequence or lacks structural variants or divergent loci present in field isolates, leading to misalignment and inaccurate expression quantifications [27]. In addition, incomplete or inaccurate genome or transcriptome assemblies can result in significant errors in the inferred expression outcomes [28].

Despite widespread awareness of these challenges, the specific impact of the selected reference genome on scRNAseq mapping behaviour in malaria field isolates has not been characterized.

In this study, we systematically evaluated how a generic versus a sample-matched reference genome sequence influences scRNAseq mapping behaviour and expression inference. We generated scRNAseq data from the circulating asexual and sexual stages, and a reference genome from the asexual stages, of a *P. falciparum* natural infection from Mali (ML52). The ML52 assembly showed high contiguity (N50 1.34 Mb), with the majority of sequences assigned to 14 chromosomes, and ∼1.35 Mb in unplaced contigs enriched for subtelomeric and repetitive regions. These features likely reflect sequence divergence, structural complexity, and mixed-strain composition. The ML52 sample was identified to have five different strains present although a single strain was dominant among the asexual stages. PfML52 isolate scRNAseq short reads were mapped against this isolate’s assembled genome and the Pf3D7 reference genome.

Our overall aim was to disentangle true biological signals from mapping- and annotation-related artefacts, establish a genome-aware framework for interpreting single-cell transcriptomic data in genetically diverse malaria parasites, and assess the utility of a matched reference genome relative to the Pf3D7 reference genome for scRNAseq analysis.

## Materials and Methods

### Sample origin and processing

The ML52 *P. falciparum* sample was collected in 2022 from a Malian symptomatic donor. The patient, a 14-year-old female, presented to the clinic in Faladie, Mali with malaria symptoms including fever (38.2°C), headache and abdominal pain. She had a parasitaemia of 13,960 parasites per µl as estimated by microscopy. A 5 mL blood draw was performed, followed by prompt treatment, and this was processed as described in [25]. Briefly, venous blood was collected in CPDA tubes and processed to separate sexual and asexual *Plasmodium* stages. Gametocytes were enriched by resuspending 3 mL of blood in a suspended animation (SA) buffer and passing it through a MACS column, followed by elution and concentration for 10x Genomics loading. The MACS flowthrough was filtered (Plasmodipur™) to remove leukocytes, then treated with streptolysin O (activated with DTT) to lyse uninfected RBCs and enrich circulating asexual-stage–infected RBCs. Cell counts were determined by hemocytometer, and parasitemia was assessed by Giemsa smear. Sexual and asexual fractions were pooled for a 10x Chromium Single Cell 3′ v3 run. The remaining fractions were stored at -80°C for high molecular weight (HMW) DNA extraction.

HMW DNA was extracted as previously described [29]. In brief, DNA was extracted from enriched parasite fractions using Qiagen’s MagAttract HMW DNA extraction kit on a KingFisher™ Apex Automated Extraction System from Thermo Fisher Scientific, and eluted in a volume of 100 μl. HMW DNA was sheared using the Covaris g-TUBE by passing the DNA through the g-Tube twice at 3500 rpm to obtain ∼10 kb fragments required for library preparation. DNA quantity was measured with a Qubit Fluorometer using Qubit’s dsDNA High Sensitivity Assay kit.

PacBio HiFi DNA sequencing libraries were constructed according to the manufacturers’ instructions using a PacBio SMRTbell® Express Template Prep Kit 2.0 and PacBio SMRTbell® gDNA Sample Amplification according to the protocol for Ultra Low Input (ULI) material. The sample was multiplexed on a single Revio flow cell plexed together with 24 other samples

### Single-cell data QC: filtering of doublets and low quality cells

#### Single-cell 10X encapsulation

Illumina short-read single-cell transcriptomes were previously generated from the enriched asexual and sexual fractions using the 10x Genomics Chromium platform as described in [25]. In brief, the protocol enables the encapsulation of cells into droplets where they are uniquely barcoded before sequencing. A majority of the droplets will not contain ‘actual’ cells but will still contain ambient RNA emanating from lysed cells that will also end up getting barcoded and sequenced and it is important to discriminate these ‘empty’ droplets/barcodes from those with ‘actual’ cells. To obtain high quality cells for this dataset, rigorous QC was performed similar to [25].

#### Cell mapping and cell calling

In summary, Cell Ranger (v9.0.1) [30] was used for mapping short reads against a combined *Homo sapiens* (GRCh38.p14) plus *P. falciparum* genome, with the maximum intron size set to 5,000 bp, to generate count matrices from the Illumina fastq sequences. Because the sample was blood-derived infected erythrocytes, it contained RNA from both parasite (*P. falciparum*) and the host (*H. sapiens*). Some uninfected host cells may have also ended up getting sequenced since the enrichment of infected erythrocytes is not 100% efficient. To retrieve ‘actual’ cells we used CellBender v0.3.2 [31] which estimates the ambient RNA profile for every cell from the raw cell ranger matrix and uses this to discriminate droplets with ‘actual’ cells from those that are ‘empty’. CellBender has been shown to capture low expression level cells such as early rings as opposed to Cell Ranger which was used in [25] which may mistake these droplets as only containing ambient RNA. Since we applied CellBender to a combined *Plasmodium+human* reference we retained only those ‘actual’ cells with at least 100 *Plasmodium* reads to increase the likelihood of capturing *Plasmodium* infected RBCs and discard barcodes containing human cells. This pre-QC CellBender-called set of *Plasmodium* cells was then subjected to stage assignment, strain assignment, and QC as discussed below.

### Stage assignment and removal of poor quality cells

We assigned stages to the pre-QC CellBender-called *Plasmodium* cells using singleR [32] with two reference atlas datasets [25]; [33]. This is different from the analysis in [25]. We used singleR here since it assigns labels to each cell using markers identified in the reference atlas and not affected by the presence of poor quality cells allowing stage assignment to pre-QC cells. This allowed the identification of lifecycle stages being removed at different parts of the QC steps to avoid biased removal of a particular stage. We then removed some cells based on CellBender estimated metrics for each cell including cells with >50% ambient RNA proportion and those with <0.9 probability of being ‘actual’ cells as estimated by CellBender. Further QC was performed on these cells to remove cells above the 99.5th percentile for mitochondrial read percentage, as well as cells below the 1st percentile or above the 99th percentile for gene count and UMI count. We identified and removed doublets using scDblfinder [34]. Any cells that were not confidently assigned to stages were also removed.

### Strain assignment and genotype-calling

We assigned strains to the pre-QC CellBender-called *Plasmodium* cells using souporcell v2.5 [35] pipeline with hisat2 v2.1.0 [36] as the aligner. We used the elbow plots to estimate the most likely K and evaluated 3 candidates for K by calculating identity-by-state IBS using the VCF and *vart*rix1.1.22 (10x Genomics) allele counts generated by souporcell [37]. This is different from the approach used in [25] where we used the minimap2 aligner and a strain-level pseudobulk re-genotyping approach to leverage the power of having more cells genotyped for each strain. The approach here allowed faster and easier processing due to the elimination of the pseudobulk re-genotyping while maintaining accuracy since hisat2 restricts mapping of transcripts to protein coding regions which are less prone to error. We filtered genotypes at two levels. First we removed loci from the VCF generated by mapping all reads in the dataset to the reference. We removed loci that had a freebayes mapping quality of < 20 and depth < 5, those that were absent in the ‘PASS’ biallelic set in the pf7 malariagen [38] dataset and those outside the nuclear genome. To account for the fact that different strains may contain variable stage representation and therefore different expression profiles and variable genotype confidence along the genome, we performed a second layer of filtering at the strain level. Here, we removed genotypes for specific strains using the vartrix allele counts per cell generated as part of the souporcell pipeline. We aggregated these allele counts across all cells in each strain into a pseudobulk metric representation of the alleles. Any loci in a strain cluster supported by less than five reads/UMI was removed as having low support. Any loci with >0.8 alternate allele fraction were called as alternate and those with <0.2 as reference with everything else as ‘low purity’. These genotypes were used to estimate IBS between pairwise strains comparisons using SNP relate [39]. The correct K was identified when further increase in K resulted in no new genotypically distinct clusters. After identifying the correct K we removed any strain doublets and any cells that were not assigned strains.

### Identification of dominant asexual strains and genotype comparison with HMW Pacbio DNA

We then used the strain assignments together with the stages assigned above to identify the most dominant asexual strain for analysis in the rest of the paper. We used the barcodes for the QCd cells in the dominant asexual strain and that from other strains to get clean genotypes from the souporcell vartrix file for each strain using the process described in the above section ‘*Strain assignment and genotype-calling*’. We then compared the genotypes from each strain to that of the PfML52 HMW PacBio genome which was generated by mapping the assembly/reads to the reference genome using bcftools mpileup (--min-MQ 30 --min-BQ 30 --skip-indels -d 5000) and call (-mv –ploidy 1) and further filtering to retain high quality biallelic SNPs with depth > 10. We then used SNPRelate IBS to compare the genotypes from all the strains in PfML52 scRNAseq data including the most dominant asexual strain with that of the PfML52 HMW DNA. Since PfML52 was generated from DNA we expected high IBS between the most dominant asexual strain and PfML52 DNA.

Figure S1 summarises the Workflow of the PfML52 natural infection QC process.

### Genome assembly

PacBio HiFi reads were mapped to the human reference genome (GRCh38.p14) using minimap2 (v2.28) [40] with the option ‘-ax map-hifi’. samtools [41] was used to extract unmapped reads with the ‘-bf 0×4’ flag. Hifiasm (v0.19) [42], a haplotype-resolved assembler optimized for high-accuracy long reads, was employed to assemble the unmapped PacBio HiFi reads with -l0 parameter to disable purging of haplotypic duplication. This generated a high-quality ML52 draft genome. The resulting contigs were scaffolded with RagTag (v2.1.0) [43], using 3D7 reference genome as a guide. MitoHiFi was used to identify mitochondrial and apicoplast genomes [44]. The coordinates and size of telomeric repeats in the scaffolded chromosomes and unplaced contigs was inferred by telofinder (v.1.0.0) [45] using the consensus sequence TT(T/C)AGGG.

### Gene annotation, orthology, and gene family assignment

Primary annotation of the PfML52 assembly was conducted using Companion [46], which transfers gene models from the Pf3D7 reference genome (PlasmoDB, v68) based on sequence similarity and conserved synteny. The Companion tool was run on the assembly without “pseudochrome contiguation” parameter, to disable additional scaffolding of the unplaced contigs, and the following parameters: “No, do not use reference protein evidence”, “Yes, perform pseudogene detection”, “Yes, use BRAKER for structural annotation”, “No, use Liftoff to transfer reference gene models”. Companion integrates multiple annotation strategies including Liftoff-based ortholog transfer, RATT-based annotation transfer, and homology-assisted gene prediction using AUGUSTUS and GeneMark. Liftoff implementation of the tool was used to map Pf3D7 gene models onto the PfML52 assembly preserving exon–intron structures and genomic collinearity. All genes that were successfully mapped inherited gene identifiers and family assignments from the corresponding Pf3D7 annotations. For loci where direct transfer was incomplete or unsuccessful due to structural variation or sequence divergence, Companion reconstructed gene models using TxRATT [47] and TxAUGUSTUS/GeneMark [48], [49] reference-guided annotation approaches while retaining association with Pf3D7 orthologs where possible. Aragorn [50] and Infernal [51] implementations were used to annotate non-coding genes (tRNA, rRNA, snoRNA, snRNA, ncRNA). For genomic regions lacking clear transferable Pf3D7 orthologs, coding sequences were predicted using the AUGUSTUS and GeneMark implementations within Companion. These predictions lack corresponding PF3D7 identifiers and include both hypothetical genes lacking clear Pf3D7 counterparts and homology-supported predictions associated with Pf3D7-derived evidence. Genes annotated by Liftoff have sequence identity and coverage values associated with them in reference to the Pf3D7 annotation. Gene coverage was defined as the fraction of the annotated Pf3D7 gene body (exonic regions) aligned to the corresponding PfML52 locus based on Liftoff orthology mappings. Gene sequence identity was calculated as the proportion of matching nucleotides across the aligned exon/CDS regions between PfML52 and Pf3D7 orthologous loci. Both metrics ranged from 0 to 1, with values approaching 1 indicating near-complete structural conservation and high nucleotide similarity between references.

Ortholog group assignments were obtained from PlasmoDB/VEuPathDB, where orthology relationships are defined using the OrthoMCL framework [52]. Single copy orthologs (SCO) represent evolutionarily conserved loci typically associated with essential cellular functions. SCO in this study are defined as orthologous genes that are present as one copy in each of six *Plasmodium* species annotated reference genomes (*P. ovale curtisi PocGH01, P. ovale wallikeri PowCR01, P. vivax PvP01, P. malariae PmUG01, P. knowlesi PKNH, and P. falciparum Pf3D7, PlasmoDB v68)*. This set was generated using the ortholog groupIDs for each gene from OrthoMCL and subsetting to those groups with only one gene in each of the above species. The rest of the genes are categorised into non single copy orthologs (non-SCO) which comprised orthologous groups in which one or more species contained two or more paralogous copies of non-multigene families and multigene families (which include *var, rifin, stevor, phist, etramp,* and *exp* genes). Multigene families are known for extensive copy number variation and structural complexity and they are frequently associated with adaptive processes such as host–parasite interactions, cytoadhesion and immune evasion.

PfML52 genes were classified into three categories: (i) Liftoff single copy genes, (ii) Liftoff non single-copy genes, and (iii) Non-Liftoff genes. Categories i and ii comprise genes where Pf3D7 gene models were successfully transferred onto the PfML52 assembly, and have corresponding Pf3D7 gene identifiers as well as the alignment coverage and sequence identity values with respect to the Pf3D7 child features (exon/CDS). Category iii comprises genes not directly transferred by Liftoff, including both reference-guided reconstructed annotations (TxRATT and TxAUGUSTUS/GeneMark) and *ab initio* gene predictions generated using AUGUSTUS and GeneMark within Companion. Some of these loci retain association with Pf3D7 orthologs through homology-supported annotation evidence, whereas others represent hypothetical genes lacking clear Pf3D7 counterparts or functional annotation. Although orthology relationships were retained where supported, all predicted genes were assigned PfML52 locus identifiers in the final annotation set. All quantification analyses were stratified according to the above categories to address differences in mapping behaviour, annotation confidence, and structural complexity.

### Read mapping and expression inferences

Single-cell RNA-seq short reads (10x Genomics, Illumina) were first used for all UMI-based gene expression quantification analyses. These short reads were independently mapped to the ML52 assembled genome and 3D7 reference genome using Cell Ranger (v9.0.1).

UMI counts were aggregated per gene across cells, and metrics such as coverage, sequence identity, and mapping quality (MAPQ) were evaluated at the chromosome and gene levels. Mapping quality scores were interpreted following STAR conventions (MAPQ = 255 for unique mapping; 3 for multi-mapping with the best alignment; 1 for low-confidence multi-mapping; and 0 denoting unmapped or ambiguous reads). However, in repetitive or paralogous genomic regions, particularly within multigene families, MAPQ = 255 does not necessarily guarantee biological uniqueness because highly similar loci may remain insufficiently resolved by short-read alignments.

### Long-read genomic alignments and structural analyses

PfML52 PacBio HiFi genomic reads generated on the Sequel II platform (Pacific Biosciences HiFi sequencing) [53] were mapped to both the PfML52 assembly and the Pf3D7 reference genome. These alignments were used to assess assembly continuity, investigate the placement of unplaced contigs, evaluate sequence divergence between the isolate and reference genomes, and provide structural context for loci exhibiting reference-dependent differences in scRNAseq quantification. Whole-genome alignments were visualized using genome dot plots generated in D-GENIES (v1.5.0) [54], while selected genomic regions were examined in IGV (v2.4.1) [55].

### Locus-level investigation of reference-dependent quantification differences

Genes showing evidence of reference-dependent differences in expression quantification were further investigated using short-read scRNAseq alignments. Mappings to the PfML52 and Pf3D7 genomes were manually inspected in IGV (v2.4.1) to compare coverage profiles, mismatch patterns, soft-clipping events, alignment positions, and annotation structures. Where relevant, observations from short-read alignments were interpreted alongside PacBio HiFi genomic alignments to distinguish mapping artefacts, annotation-related effects, and underlying genomic differences.

### Gene-level confidence assessment

To evaluate scRNAseq data mapping confidence, annotation accuracy was factored by taking into account the coverage (“coverage”) and sequence identity (“sequence_Id”) values provided by Liftoff implementation in Companion. Moreover, only uniquely mapped reads (MAPQ = 255) were quantified. These metrics were used to assess the completeness and accuracy of mappings to both PfML52 and Pf3D7 genomes. Genes retained for downstream analyses met stringent coverage criteria (0.98–1.0). Whereas sequence identity was evaluated across its full observed range.

### Differential expression inferences assessment

Difference in expression inference between the same scRNAseq data mapped to each of the two reference genomes was determined using gene-level unique molecular identifier (UMI) counts aggregated across all single cells which passed quality control filters. Genes were considered differentially quantified dependent on the reference genome if they showed: a fold-change greater than 1.2, with total sum-UMI count exceeding 100 (UMI_sum_per_gene = UMI_inPf3D7 + UMI_inPfML52), statistical significance defined by both Poisson test p-value and Benjamini–Hochberg adjusted p-value (padj < 0.05). These criteria were applied consistently both before and after the exclusion of unplaced contigs in the ML52 genome. Comparisons were done for genes assigned to chromosomes only and those on unplaced contigs, and independently for single copy and non single-copy loci as previously defined, to account for any differences in annotation ambiguity.

### Multigene families analysis and reference-dependent mapping behaviour

We performed a dedicated analysis of multigene families, including *var, rifin, stevor, phist, etramp, export, rpb,* and *sica* genes. Genes within each multigene family were classified into three categories: (i) PfML52 genes with a corresponding Pf3D7 gene ID, referred to as shared genes, (ii) PfML52 genes lacking a corresponding Pf3D7 gene ID, and (iii) Pf3D7 gene identifiers not present in the PfML52 Companion output. Expression inferences were initially assessed using raw UMI-based heatmaps for shared and non-shared genes in all the abovementioned multigene families. For heatmaps, quantification values were aggregated by developmental stage (early ring, late ring, early trophozoite, developing gametocyte, canonical female gametocyte, non-canonical female gametocyte, and male gametocyte) and compared between mappings to the sample genome and the reference genome.

Reads were classified according to their alignment behaviour across references. Reads aligning to annotated *var* genes in both the PfML52 and Pf3D7 mappings were designated as shared reads. Here, shared refers to the same sequencing read being aligned in both references and does not imply alignment to orthologous genes or to the same gene. Reads aligning only to PfML52 *var* genes were classified as PfML52-specific, whereas reads aligning only to Pf3D7 *var* genes were classified as Pf3D7-specific. Then, to determine whether *var* shared reads mapped to orthologous loci or were redistributed among paralogous genes, PfML52 *var* genes with more than 500 shared reads were examined with bipartite gene–gene matrices and heatmap visualisation.

For further validation, the top *var* genes in each comparison with the highest number of reads were selected, and read redistribution was visualised using alluvial plots. Additionally, Short-read coverage profiles for the top three *var* genes with highest read counts in each comparison were manually inspected in the IGV genome browser.

## Results & Discussion

Here we investigated whether the choice of reference genome impacts scRNAseq expression inference in *P. falciparum*. We integrated a new isolate-specific genome assembly with corresponding scRNAseq data from the same natural malaria infection and compared these results to when the same scRNAseq data are mapped to Pf3D7 to evaluate how the selected reference genome influences short-read mapping behaviour and expression inference.

### PfML52 genome assembly and annotation

Assembly of PacBio HiFI reads of PfML52 by hifiasm resulted in 75 contigs, with an N50 of 1.34 Mb (megabase), comparable to the theoretical N50 of ∼1.688 Mb for Pf3D7, corresponding to chromosome 10, indicating high-contiguity. The largest contig spanning 3.23 Mb is similar in size to the largest Pf3D7 chromosome 14 (3.29 Mb) (Fig. 1A, Fig. 2A,B). MitoHiFi [44] identified mitochondrial and apicoplast genomes amongst 2 of these contigs. Scaffolding by RagTag [43] assigned 26 contigs onto 14 chromosomes (total 23.47 Mb), with 12 gaps, whereas 47 contigs (1.35 Mb) remained unplaced. Upon further examination, these unplaced contigs mapped well to regions of the ML52 genome (Fig. 1B), and thus are likely to comprise additional allelic variation from the natural infection, which did carry multiple strains .

**Figure 1.**
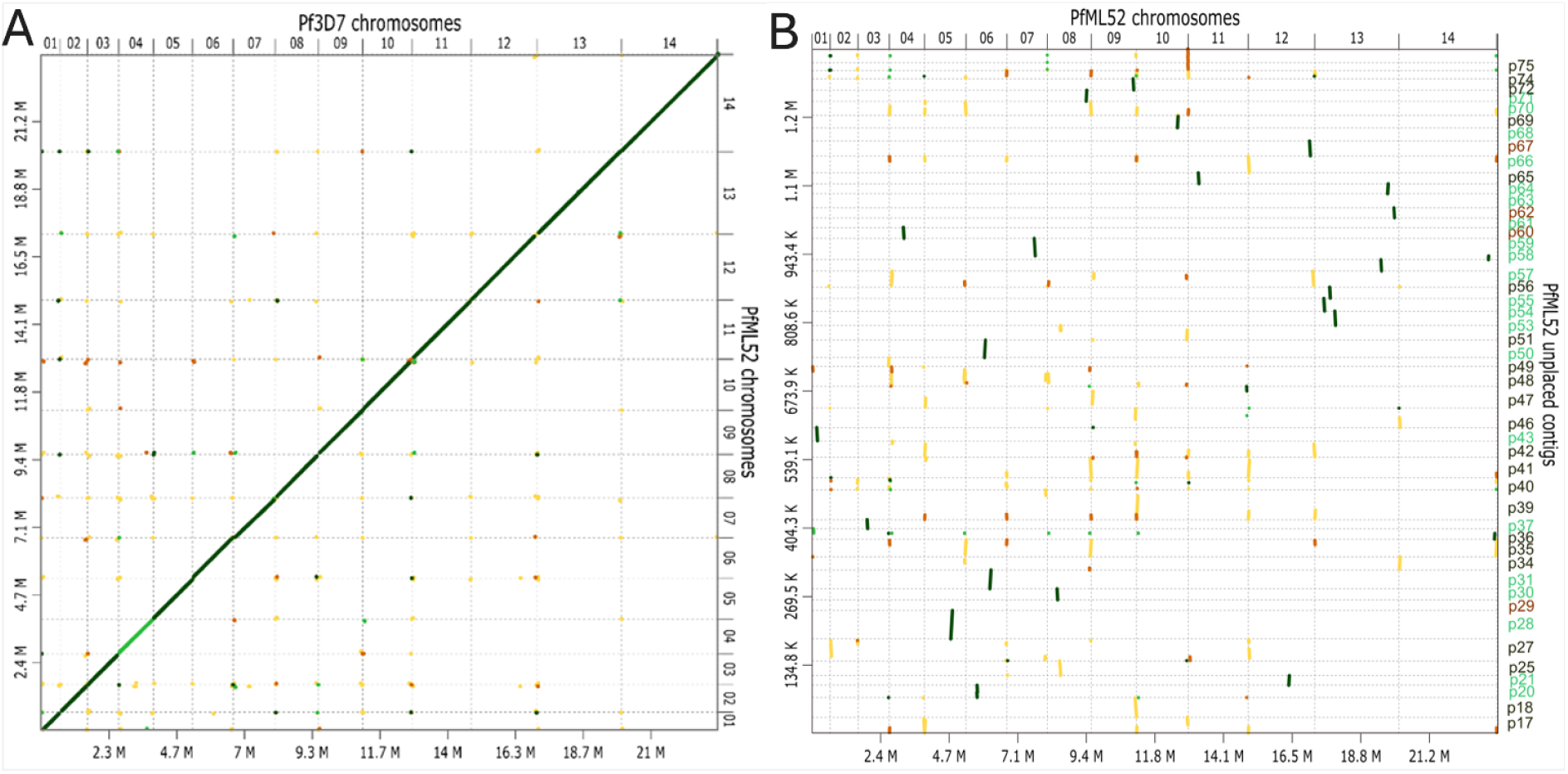
Genome alignment dot plots of PfML52 and Pf3D7 genomes. A. PfML52 genome assembly shows high global collinearity with the Pf3D7 reference genome. B. Genome dot plot comparing PfML52 unplaced contigs against its chromosomal assembly, illustrating partial alignments of unplaced contigs to subtelomeric and repetitive chromosomal regions.

**Figure 2.**
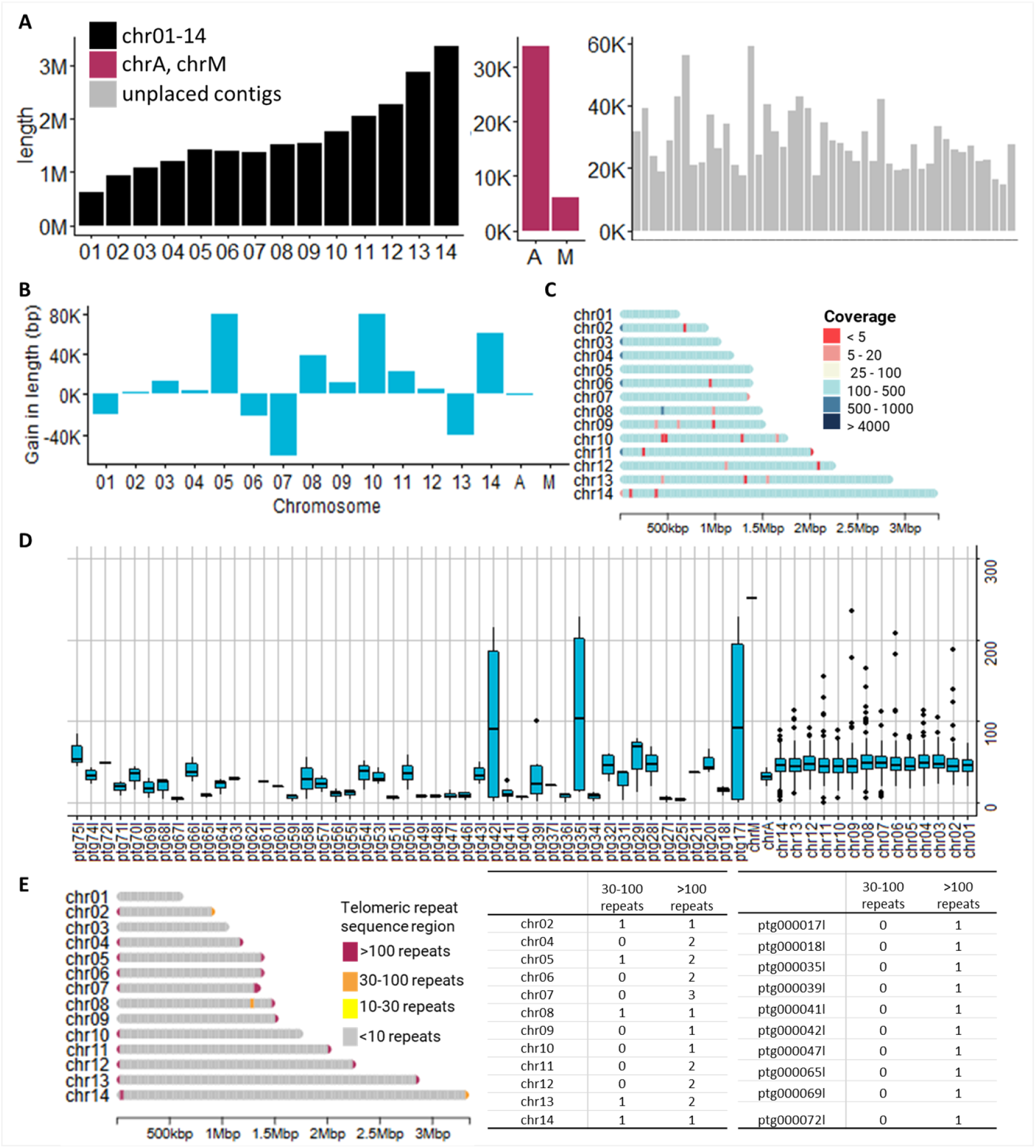
Structural features, coverage, and telomeric repeat distribution in the PfML52 genome assembly. A. Length of contigs in the final scaffolded assembly. B. Gain in length for each chromosome of the assembly compared to the Pf3D7 reference. C. Coverage across each chromosome of the scaffolded assembly, in bins of 10,000 bp regions and colored according to the values denoted in the legend. D. Coverages across the genome for each chromosome and unplaced contig displayed as boxplots of 10,000 bp regions. E. Telomeric repeat regions were inferred using telofinder across the genome of PfML52, and displayed here for the chromosomes, colored by the number of repeats in 10,000 bp bins. The table on the right shows the number of such bins with 30-100 or >100 repeats across the chromosomes and unplaced contigs.

These unplaced contigs ranged from 14,800 to 58,822 bp and could not be confidently assigned to chromosomes during scaffolding (Fig. 2A). PfML52 is a clinical sample estimated to contain 5 strains with a single major dominant strain. Therefore, unplaced contigs may reflect divergent haplotypes and/or unresolved repetitive subtelomeric sequence already partially represented within the primary chromosomal assembly. Of the 47 unplaced contigs, 21 contigs aligned to the core genome, 22 aligned to the subtelomeric ends of the genome and 4 were unmapped (Fig. 1B).

In the chromosomal assembly, the 14 chromosomes have median depth of 48x, with very few low coverage regions, amounting to ∼3% of 10 kbp bins covered by < 25 reads (Fig. 2C,D). Telomeric repeat sequences (GGGTT[T/C]A) [56] were identified at 21 chromosomal ends out of the expected 28, emphasizing the completeness of the ML52 assembly (Fig. 2E). 10 of the unplaced contigs harbor these repeats as well and likely constitute the remnant chromosomal ends (Fig. 2E). Overall, the PfML52 assembly displayed high sequence similarity and synteny to Pf3D7, suggesting a highly conserved genome organisation.

The annotation results further support this finding. Most genes from Pf3D7 were successfully transferred to PfML52, with high sequence coverage and sequence identity, suggesting good amino acid and nucleotide sequence conservation across isolates. In total 6,770 genes were annotated (6,892 transcripts) in the PfML52 genome compared to the 5,797 genes in the Pf3D7 genome (Fig.3, Table. 1). 5,262 genes from Pf3D7 were directly transferred demonstrating that the majority of Pf3D7 genes are reliably annotated via sequence similarity and orthology. The additional 1,508 annotations were derived from *ab initio* gene predictions by Companion implementations of AUGUSTUS and GeneMark as well as non-coding gene predictions by Aragorn and Infernal (Table. 1). Unplaced contigs are overrepresented in genes annotated as “hypothetical protein” comprising ∼62% (395/630), with the four contigs previously referred to as unmapped comprising entirely of them. Multigene family genes such as *rifins*, *vars*, *stevors*, etc comprised ∼14%. 42 of the protein coding genes of Pf3D7 had no apparent corresponding gene identifier in PfML52, with 35 of these belonging to multigene families. Of the rest, 5 were ribosomal genes in the apicoplast and 2 genes, *PF3D7_1366900* and *PF3D7_1366700*, correspond to an assembly gap in ML52 (chr13:2,693,757-2,693,856) leading to lack of these annotations in ML52. On the other hand, 1,202 annotations in PfML52 derived from the ab initio tools in Companion (AUGUSTUS, INFERNAL, Aragorn, Genemark) had no corresponding Pf3D7 gene identifiers (Table. 2). A majority of these are belonging to multigene families and genes labelled as “hypothetical protein” or “hypothetical protein, conserved”, with no putative functional domains. These “hypothetical” genes are present almost exclusively at the telomeric ends beyond the *vars*, and likely represent potentially false annotation (Fig. 3B).

**Figure 3.**
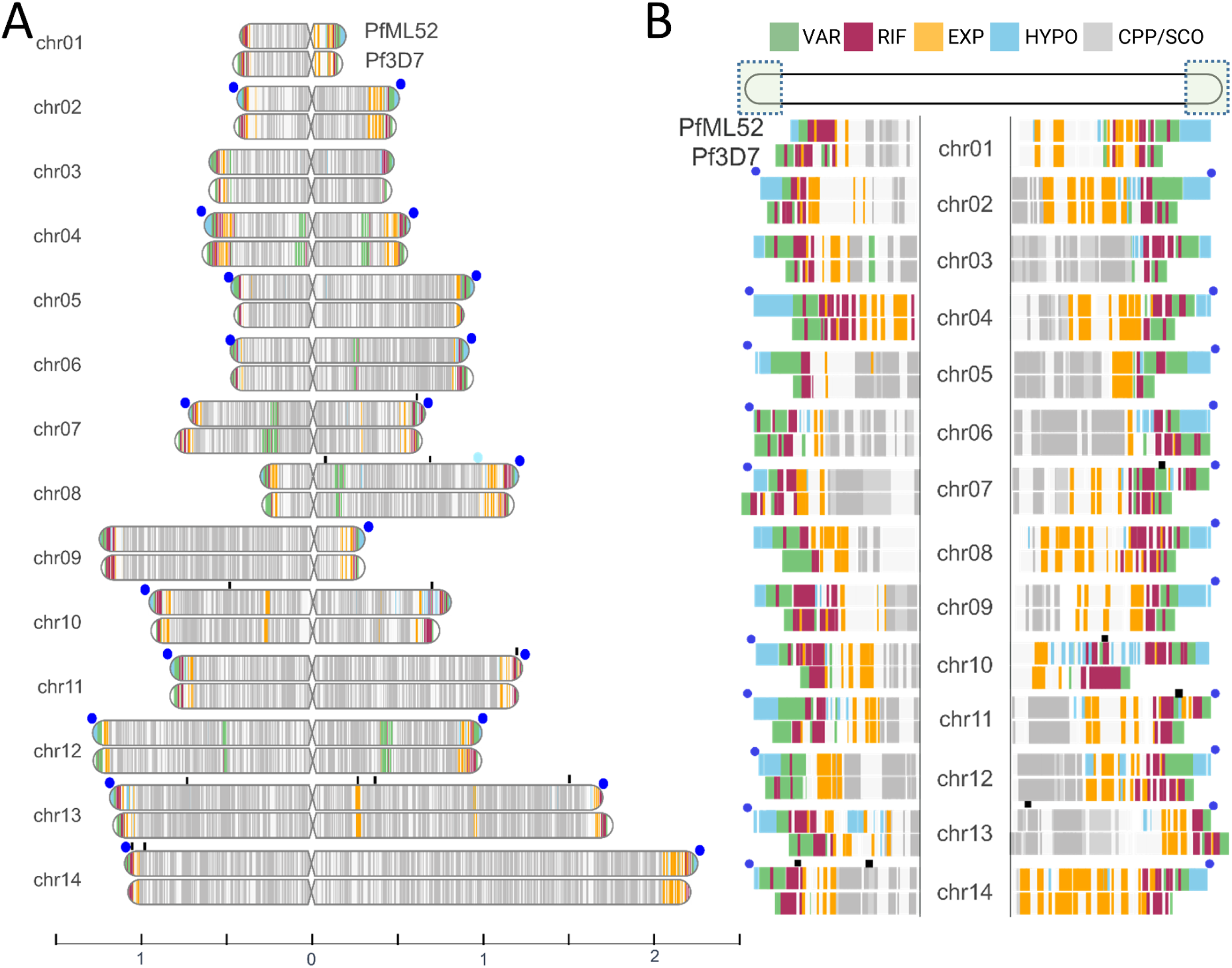
Annotation of PfML52 assembled genome. **(A).** PfML52 assembly compared to Pf3D7, by aligning along the centromeric locations, shows gene synteny across most of the genome. The chromosome scale is displayed below in Mbp. **(B).** Chromosomal ends of Pf3D7 and PfML52 show loss of gene synteny closer to the telomeric ends, with the positions of gene annotations in grey corresponding to single copy orthologs (SCO) and conserved *Plasmodium* proteins (CPP) in agreement between the two assemblies. The figure also highlights “hypothetical proteins” genes, likely false annotation, because they are present exclusively at the telomeric ends beyond the *var* genes and lack any putative functional domains. In A and B, blue circles indicate regions where telomeric repeats were identified in the ML52 genome. Black vertical marks above the chromosomes indicate assembly breaks. VAR - *var* genes, RIF - *rifin*, EXP - genes coding for the families of exported proteins (SICA, RESA, PHIST, *Plasmodium* exported protein, reticulocyte binding protein, early transcribed membrane protein), HYPO - genes labeled “hypothetical protein, ”hypothetical protein, conserved”, CPP - genes labeled ”conserved * protein, unknown function”, SCO - single copy orthologs across human infecting *Plasmodium* species.

**Table 1.** Gene IDs overlap between Pf3D7 and Companion-derived PfML52 annotation. The table lists the gene categories with protein coding PF3D7 IDs not represented in the Companion annotation (either as 1:1 annotation correspondence or through ortholog relatedness in the GeneDB column), and PfML52 IDs with no apparent Pf3D7 corresponding IDs. Values shown in parentheses represent the subset of genes exhibiting reliable expression, defined as genes with UMI counts >100 and confirmed by manual inspection in the Integrative Genomics Viewer (IGV).

| Gene category | No. of PF3D7 gene IDs not represented in PfML52 annotation | No. of gene IDs in PfML52 with no apparent PF3D7 ID |
| --- | --- | --- |
| **VAR** | 0 | 83 (5) |
| **RIFIN** | 27 (2) | 81 (2) |
| **STEVOR** | 4 | 19 |
| **SICA** | 0 | 2 |
| **Other** | 11 | 853 (8) |
| **Non-coding genes (tRNA, rRNA, snoRNA, ncRNA)** | 0 | 164 |

**Table 2.** Redistribution of reads from the three dominant PfML52 *var* genes across the Pf3D7 *var* repertoire. For each highly expressed PfML52 *var* gene, all reads assigned to that locus were compared with alignments to the Pf3D7 reference. The table reports the number of Pf3D7 *var* genes sharing reads with each PfML52 gene and the total number of shared reads, before and after applying a threshold of >500 shared reads. These values quantify read redistribution among homologous *var* loci rather than one-to-one orthologous relationships.

| Dominant PfML52 *var* genes | Without threshold |  | With threshold >500 shared reads |  |
| --- | --- | --- | --- | --- |
|  | No. Pf3D7 *var* genes sharing reads | n_shared reads | No. Pf3D7 *var* genes sharing reads | n. shared reads |
| *PfML52_000020900* | 56 | 96982 | 14 | 93394 |
| *PfML52_000340700* | 53 | 202030 | 22 | 198135 |
| *PfML52_000454300* | 57 | 43548 | 6 | 36495 |

Table S1 summarises the annotation features and annotation sources for the PfML52 genome, using the Pf3D7 reference genome.

The following table summarises the overlap in gene IDs between the Pf3D7 annotation and the Companion-derived PfML52 annotation. Candidate genes were defined as those with UMI counts >100 and subsequently validated by manual inspection in IGV.

Among the 83 var genes unique to the PfML52 annotation, five genes met these criteria (*PfML52_000215900, PfML52_000253900, PfML52_000415500, PfML52_000020900,* and *PfML52_000454300*). The latter two genes are discussed in detail in the downstream analyses. In contrast, among the 81 RIFIN genes unique to the PfML52 annotation, only *PfML52_000193100* and *PfML52_000102500* satisfied the selection criteria, whereas the remaining genes showed absent or minimal expression. Similarly, of the 27 Pf3D7 RIFIN genes without annotated PfML52 orthologues, only *PF3D7_1240700* and *PF3D7_0421500* fulfilled the same criteria. The 19 STEVOR genes unique to the PfML52 annotation showed virtually no detectable expression, with none meeting the selection criteria.

The "Other" gene category comprised the Rest, Hypothetical (Hypo), CPP, and PHIST-RESA gene groups. Among the 821 PfML52 hypothetical protein genes lacking identifiable Pf3D7 orthologues, only *PfML52_000448700*, *PfML52_000456800*, and *PfML52_000508300* fulfilled the selection criteria. The PHIST-RESA family was represented by nine genes unique to the PfML52 annotation; however, only *PfML52_000106400* and *PfML52_000394400* met the selection criteria, whereas *PfML52_000393000* was annotated as a pseudogene. Two CPP genes unique to the PfML52 annotation (*PfML52_000059400* and *PfML52_000471200*) also fulfilled the selection criteria.

### Single-cell data QC: filtering of doublets and low quality cells

After QC, and assignment of strains and stages, we obtained five strains (SC1; 3,861 cells, SC2; 6,567, SC3;1,954, SC4; 412, SC5; 2,400) spread over the asexual stages (total N = 7197) and sexual (total N = 7997) stages (Figure 4A-B). One strain, SC2, was dominant in the asexual blood stages constituting 92% of all cells in this stage (Figure 4B). The genotypes of this strain were highly concordant with those of the HMW DNA fraction with an IBS of 0.98 giving confidence that the single cell transcriptome and bulk DNA were from the same sample (Figure 4C).

**Figure 4.**
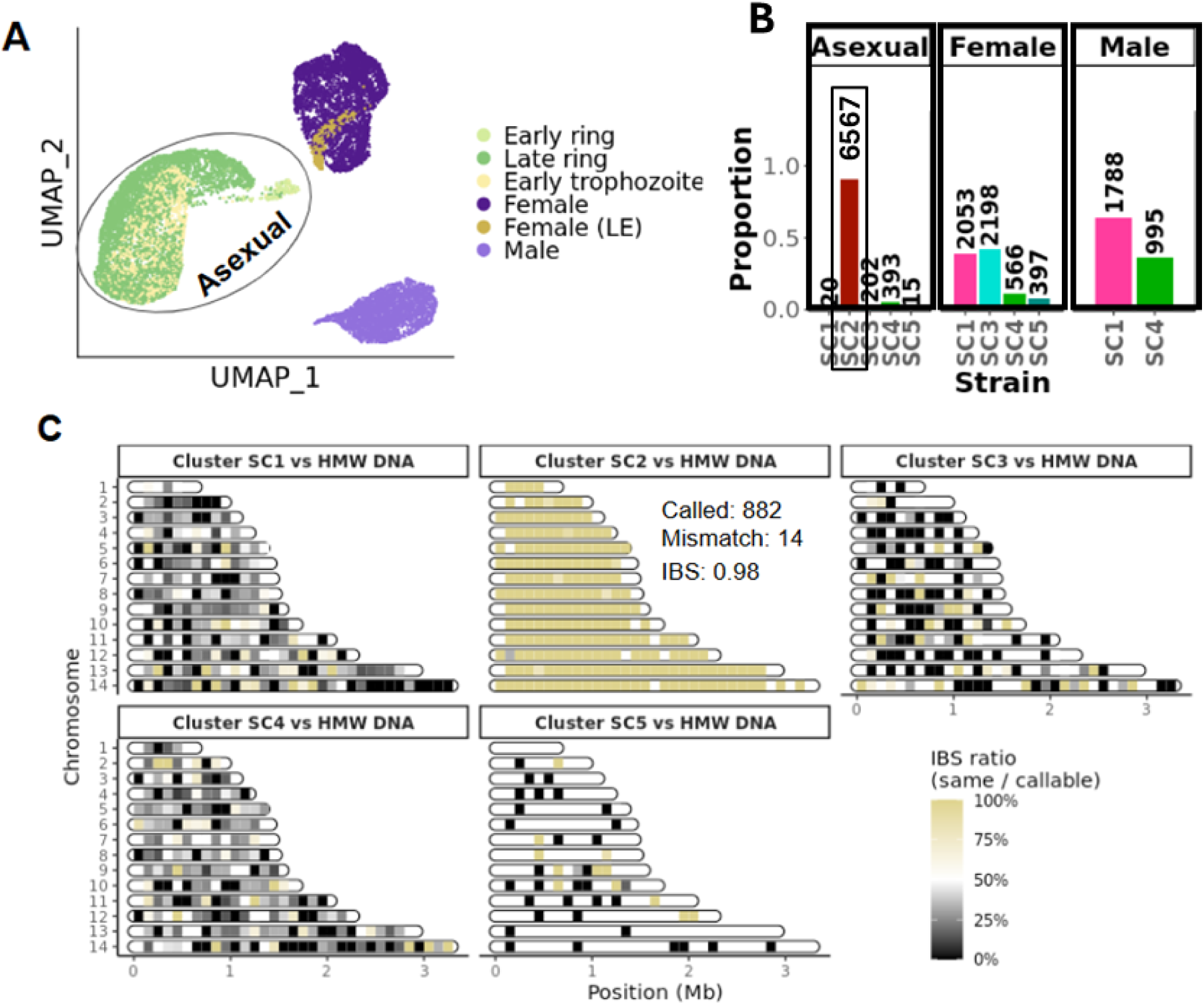
Single-cell transcriptomic profiling of *Plasmodium falciparum* natural infections across parasite strains and datasets. (A) UMAP of all parasite cells from donor PfML52 coloured by assigned stage (B) Relative proportions (Y axis) and counts (above each bar) of strains (SC1–SC5) in the asexual, female and male parasites stages. (C) Chromosome painting plot showing the comparison of overlapping SNPs between different strain clusters in PfML52 scRNAseq data and the PfML52 HMW DNA based reference genome, which was prepared from the asexual fraction. The identity between the dominant strain in the asexual stages and the DNA used to generate the assembly is expected.

### Global mapping patterns of scRNAseq short reads

Short-read scRNAseq alignments showed comparable overall mapping rates to both genomes. 48,191,488 reads mapped to PfML52 and 47,859,731 mapped to Pf3D7 from a total of 265,163,522 reads (Fig. 5A). Mapping quality distributions were also highly similar between the references with around 69% of reads mapping uniquely (MAPQ = 255), and the remainder distributed across the intermediate and low-confidence categories (Fig. 5B).

**Figure 5.**
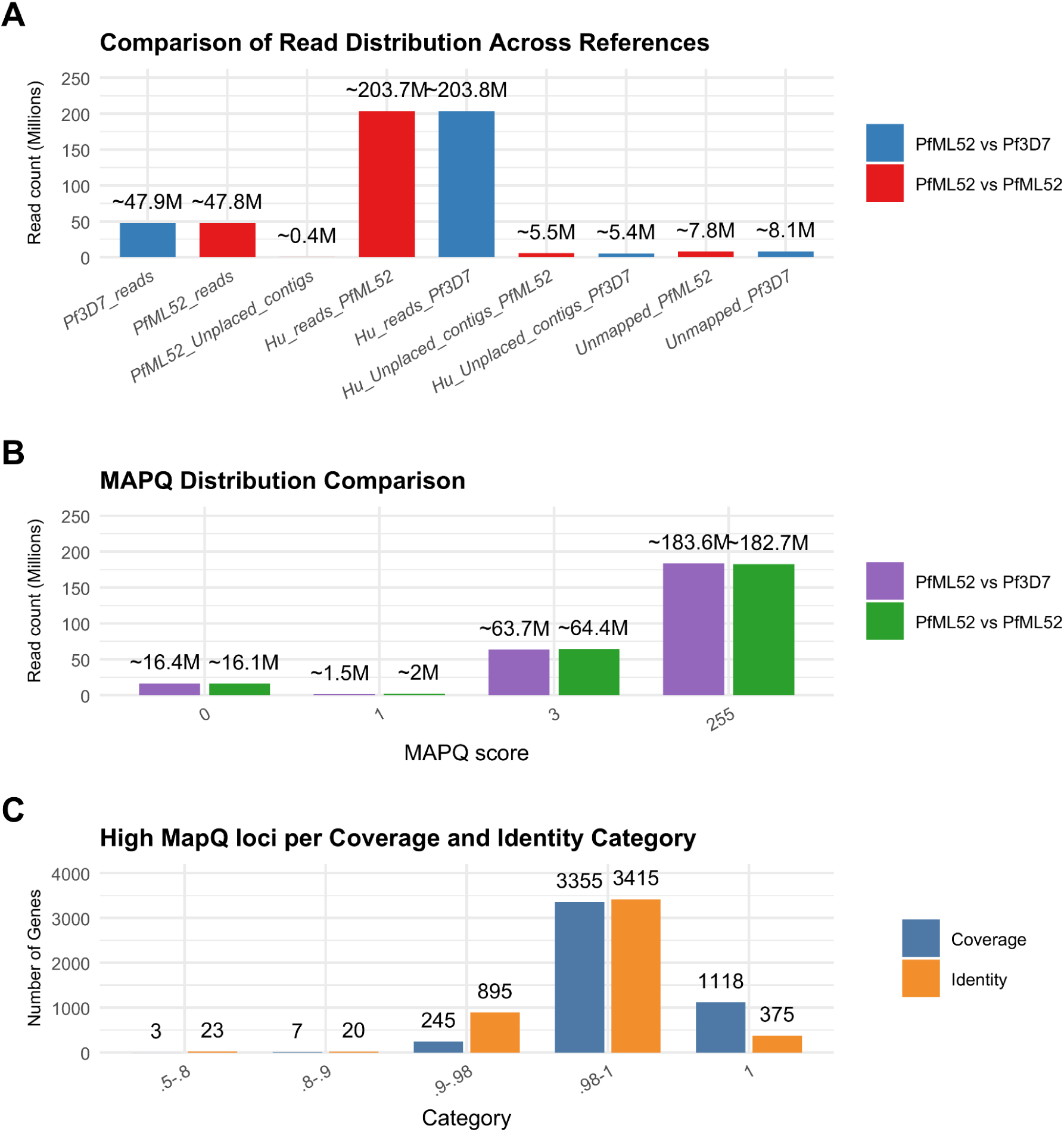
Global mapping performance and gene-level alignment metrics. (A) Total number of single-cell RNA-seq reads aligned to the ML52 isolate genome and the 3D7 reference genome, showcasing the reads distributed across parasite chromosomes, human chromosomes, unplaced contigs, and unmapped reads for PfML52 versus PfML52 and PfML52 versus Pf3D7 alignments. (B) MAPQ score distributions for both comparisons, with the majority of reads assigned MAPQ 255 and smaller fractions assigned MAPQ scores 0, 1, or 3. (C) Number of high-MAPQ loci grouped by coverage and identity categories, with counts displayed above each bar. For the first part of downstream analyses genes were required to have coverage between 0.98 to 1, whereas no threshold was applied to sequence identity.

### Quantification of UMI counts between PfML52 and Pf3D7 High MapQ genes

Among 4,728 Liftoff-derived genes in ML52 supported by uniquely mapped reads (MAPQ = 255), the majority of these were genes that showed highly similar sizes to their Pf3D7 counterparts (coverage 0.98-1 or 1 for 4,473 genes) and high sequence identity (identity 0.98-1 or 1 for 3,790 genes) (Fig. 5C). This indicates that most expressed genes are well-conserved and reliably captured, regardless of the choice of reference genome. Only reads with MAPQ= 255 are included in the first part of this study to eliminate multi-mapping artefacts. These high MAPQ genes, further categorised into 2,919 single copy orthologs (SCO) across human infecting *Plasmodium* species, and 1,809 non-single copy orthologs (non-SCO), as described in Methods section, were included in the first framework of comparison. Gene-level UMI counts showed high correlation between PfML52- and Pf3D7-based mappings. The outcomes also revealed that some genes had UMI counts higher in Pf3D7 than in PfML52 which was unexpected. These observations prompted further analysis.

To further explore why some genes had higher UMI counts when mapped to Pf3D7 than to PfML52, genes were further filtered using a more stringent coverage threshold (coverage = 0.98–1.0). Only these loci with near-complete alignment across references were retained for quantitative comparison. Differential quantification analysis was then performed using defined criteria: a fold change > 1.2 (to capture modest but systematic differences caused by reference mapping); a minimum expression threshold (UMI_sum > 100) was imposed to exclude lowly expressed genes, where stochastic sampling and UMI sparsity can inflate variability and obscure technical biases. This initial analysis resulted in 34 conserved SCO and 25 non-SCO genes showing evidence of estimated expression level being affected by choice of reference genome. Surprisingly, the majority of these genes showed higher UMI counts in Pf3D7 compared to PfML52 (Fig. 6).

**Figure 6.**
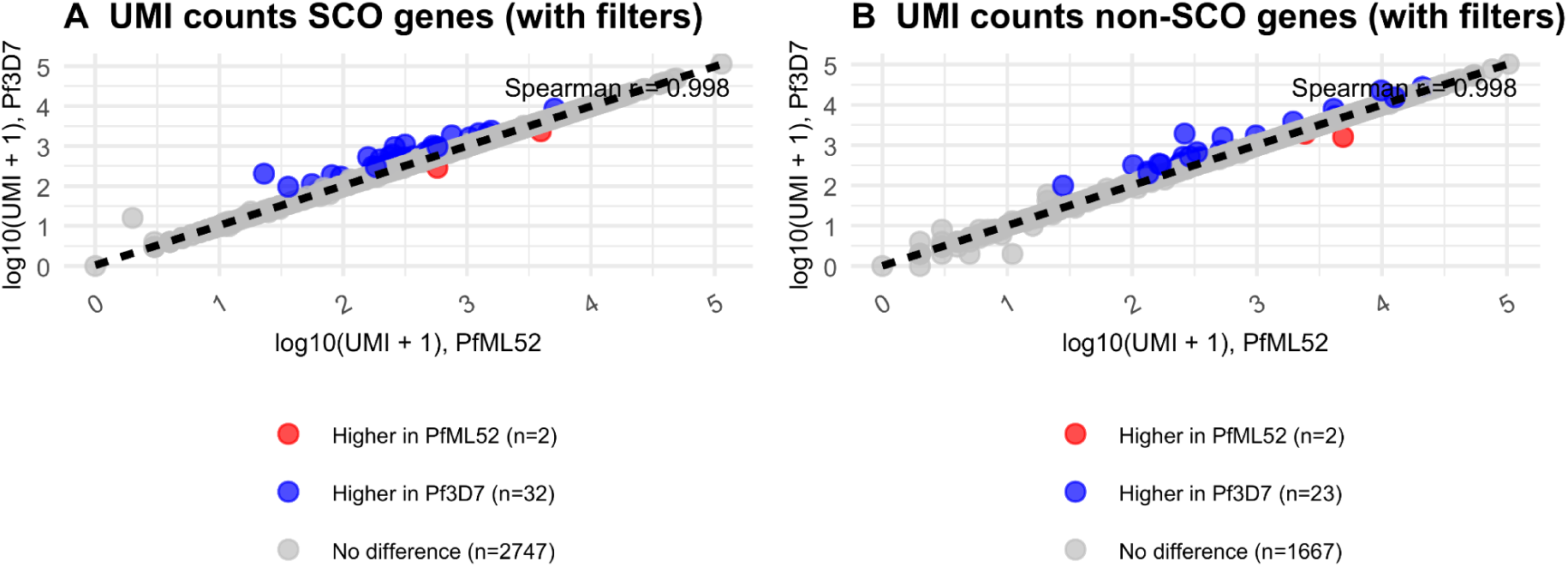
Evaluating UMI-counts concordance between PfML52 & Pf3D7 mappings and the effect of filtration across gene categories. Correlation of log10-transformed UMI counts between PfML52 and Pf3D7 stratified by gene class, showing **(A)** single-copy orthologs (SCO) and **(B)** non-single-copy orthologs (non-SCO). Genes were filtered using stringent criteria: coverage between 0.98 and 1.0, total UMI counts (UMI_sum) > 100, absolute log2-fold-change ≥ 0.263 (fold change > 1.2), and statistical significance defined by both Poisson test p-value and Benjamini–Hochberg adjusted p-value (padj < 0.05). Red and blue points indicate genes with significantly higher quantification in PfML52 and Pf3D7, respectively.

Reads mapping to regions of interest were inspected using Integrative Genomics Viewer (IGV). This inspection revealed that, in many cases where UMI counts were lower in ML52 relative to the Pf3D7 mapping, reads were partitioned between homologous loci located on ML52 chromosome contigs and corresponding regions represented on ML52 unplaced contigs. This distribution reduced the number of reads assigned to individual annotated loci, thereby artificially lowering gene-level UMI counts in the ML52 mapping (Fig. 7).

**Figure 7.**
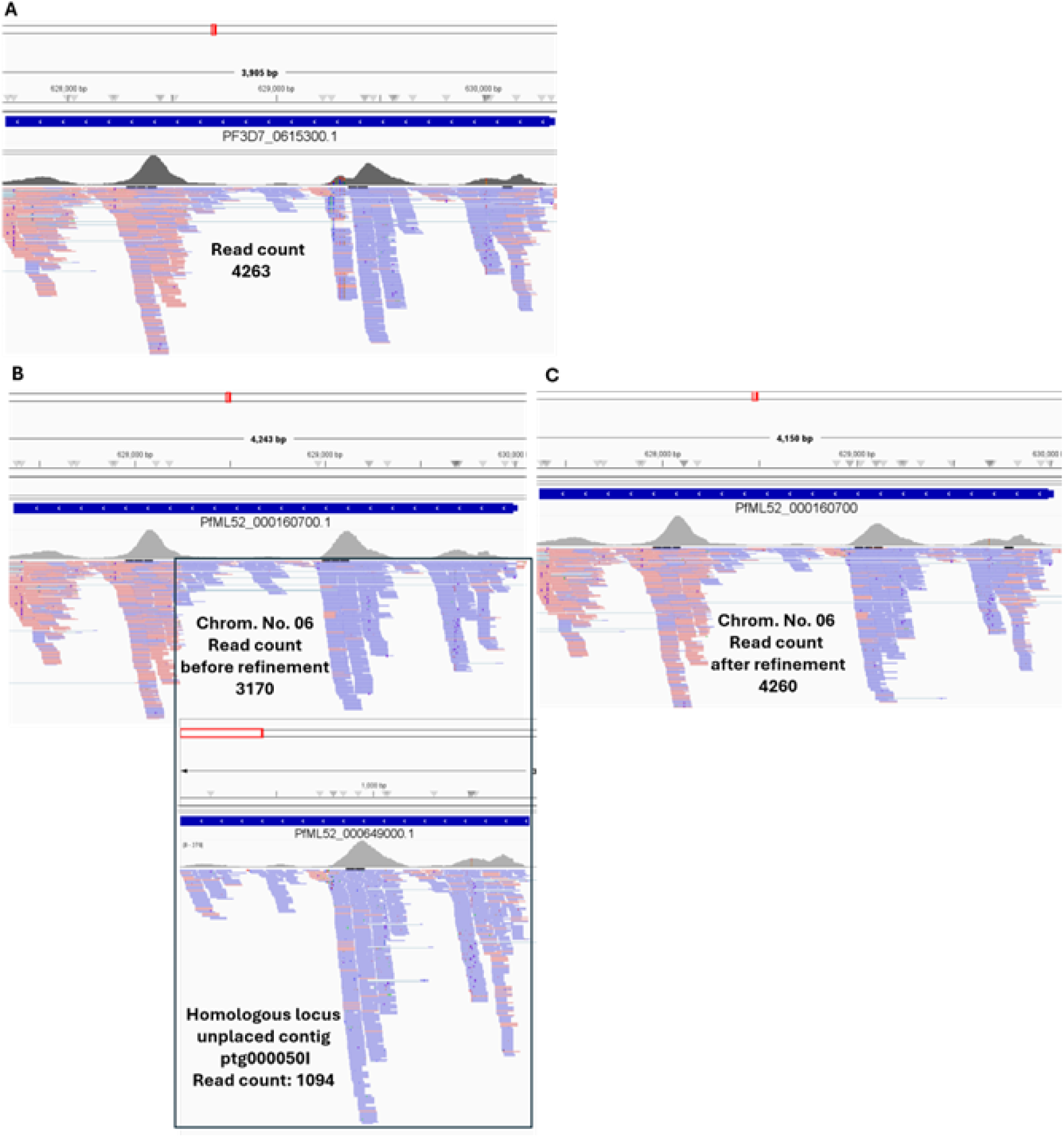
A case where reads are partitioned between homologous loci in a chromosomal contig and a corresponding unplaced contig in the PfML52 genome. **(A)** IGV view of the Pf3D7 reference showing the orthologous locus (*PF3D7_0615300*) with all reads coherently assigned, resulting in a total read count of 4263. **(B)** Corresponding PfML52 mapping illustrating read partitioning between *PfML52_000160700* on chromosome 06 (3,170 reads) and a homologous locus, *PfML52_000649000*, located on unplaced contig ptg000050I (1,094 reads). This split distribution reduces the read and UMI counts assigned to the chromosomal gene compared to its Pf3D7 ortholog. **(C)** After excluding unplaced contigs when mapping scRNAseq data to ML52, reads are consolidated at *PfML52_000160700*, increasing its read count to 4260 and restoring consistency with the Pf3D7 locus, demonstrating the impact of resolving unplaced contig-associated read partitioning.

Upon identifying that unplaced contigs in the ML52 genome were attracting some of the reads, these unplaced contigs were removed and scRNAseq alignments were recomputed against the chromosomal PfML52 assembly. This reduced the number of differentially quantified genes to 10 single-copy (SCO) and 9 non–single-copy (non-SCO) loci (Fig. 8A,B). Venn diagrams were used to visualise and compare the sets of loci that were differentially quantified before and after genome-aware mapping refinement (Fig. 8C,D).

**Figure 8.**
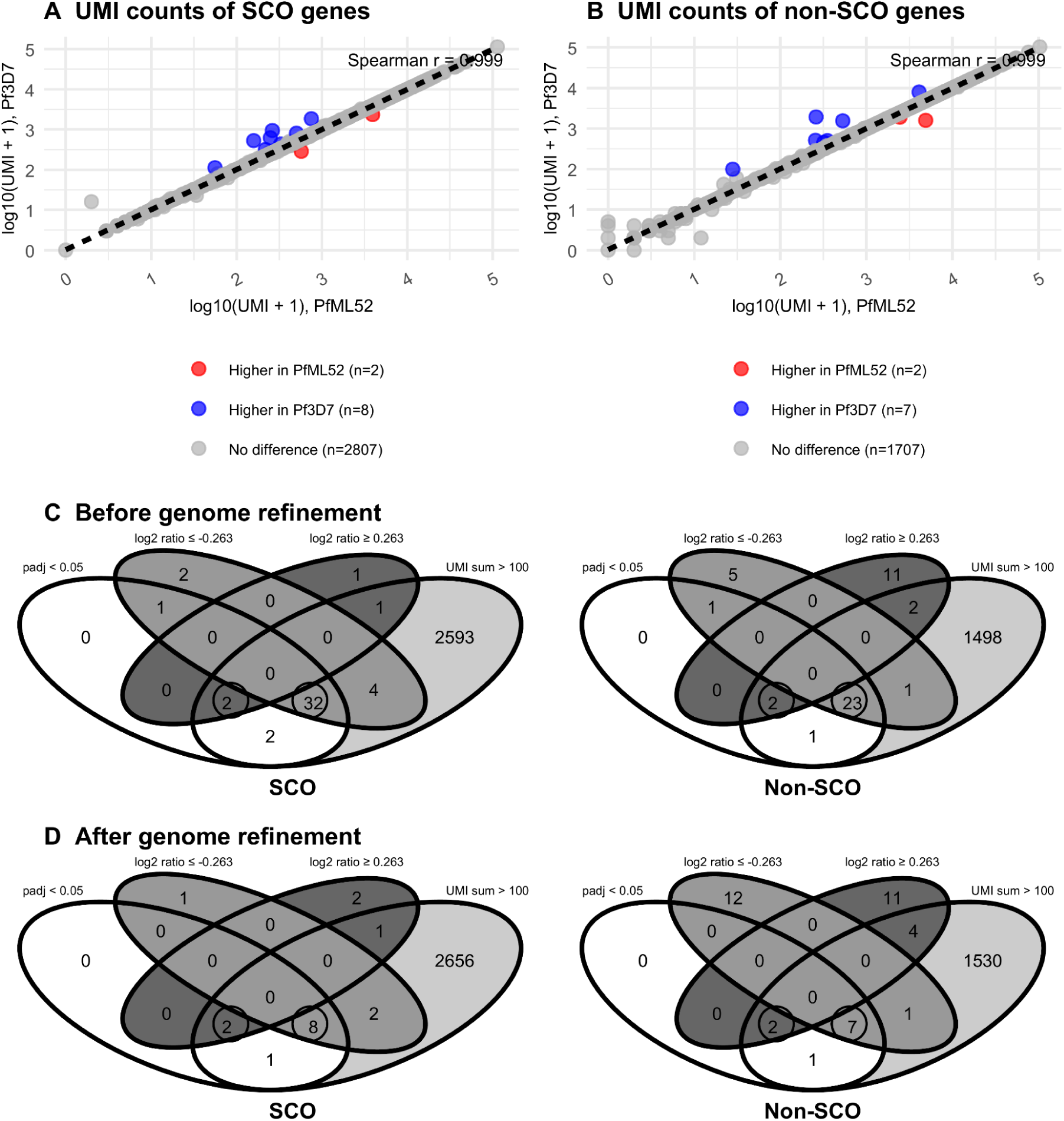
Effect of genome-aware refinement on concordance of gene quantification. **(A–B)** Correlation of UMI counts between PfML52 and Pf3D7 mappings after genome-aware refinement, shown separately for **(A)** SCO and **(B)** non-SCO orthologs. **(C–D)** Venn diagrams showing overlap of genes identified as differentially quantified (absolute log2-fold-change ≥ 0.263, UMI_sum > 100) between PfML52 and Pf3D7 mappings, stratified by SCO and non-SCO **(C)** before genome-aware refinement and **(D)** after refinement. Each circle represents the set of genes identified as differentially quantified in one reference, and overlaps indicate genes consistently detected in both mappings. Genome-aware refinement, accounting for unplaced contigs, substantially reduces the number of apparently differentially quantified genes.

Genes showing evidence of being affected by the reference genome were visually inspected in IGV. A subset of the remaining differentially quantified loci can be explained by annotation-related effects.

Specifically, twelve genes exhibit differences attributable to inconsistencies in gene model structure between the two references (supplementary data, Table S1).

In these cases, one genome contains multiple annotated isoforms or closely spaced gene models, whereas the corresponding region in the other genome is represented by a single or fewer annotations. Consequently, reads are partitioned across multiple features in the more complex annotation but aggregated into a single locus in the simpler one, producing apparent differences in UMI counts that do not reflect underlying transcriptional variation. An example gene is *PfML52_000553600 / PF3D7_1425200* (Fig. 9A). Another example is the cysteine desulfurase locus (*PfML52_000196500 / PF3D7_0716600*), where locus-level inspection revealed clear alignment inconsistencies when reads were mapped to the Pf3D7 reference. Despite being classified by STAR as uniquely mapped (MAPQ = 255), these reads exhibited fragmented alignment patterns, including mismatches, local positional shifts, and variable CIGAR structures. In contrast, mapping to the ML52 reference produced more coherent alignments, with consistent start and end positions and reduced mismatch frequency. This pattern indicates that sequence divergence between the isolate and the Pf3D7 reference introduces alignment heterogeneity, which can propagate to downstream quantification and contribute to discrepancies in inferred expression (Fig. 9B). A third example is *PF3D7_0631000 / PfML52_000176600* (tetratricopeptide repeat protein), which shows higher UMI counts in the Pf3D7 mapping relative to PfML52. Inspection of read distribution indicates that a subset of reads assigned to *PF3D7_0631000* is not recovered at the corresponding PfML52 locus. Instead, these reads preferentially align to a closely related paralogous locus in PfML52 (*PfML52_000146600*), identified as the nearest BLAST hit. As a result, reads originating from *PfML52_000176600* are partially distributed to *PfML52_000146600*. This distribution is consistent with high sequence similarity between the loci, particularly across an extended C-terminal region, and results in reduced read and UMI counts at *PfML52_000176600* relative to its Pf3D7 ortholog (Fig 9C). These findings indicate that the initial differences are primarily driven by annotation structure and assignment of reads among paralogous loci, rather than genuine differences in gene expression inference.

**Figure 9.**
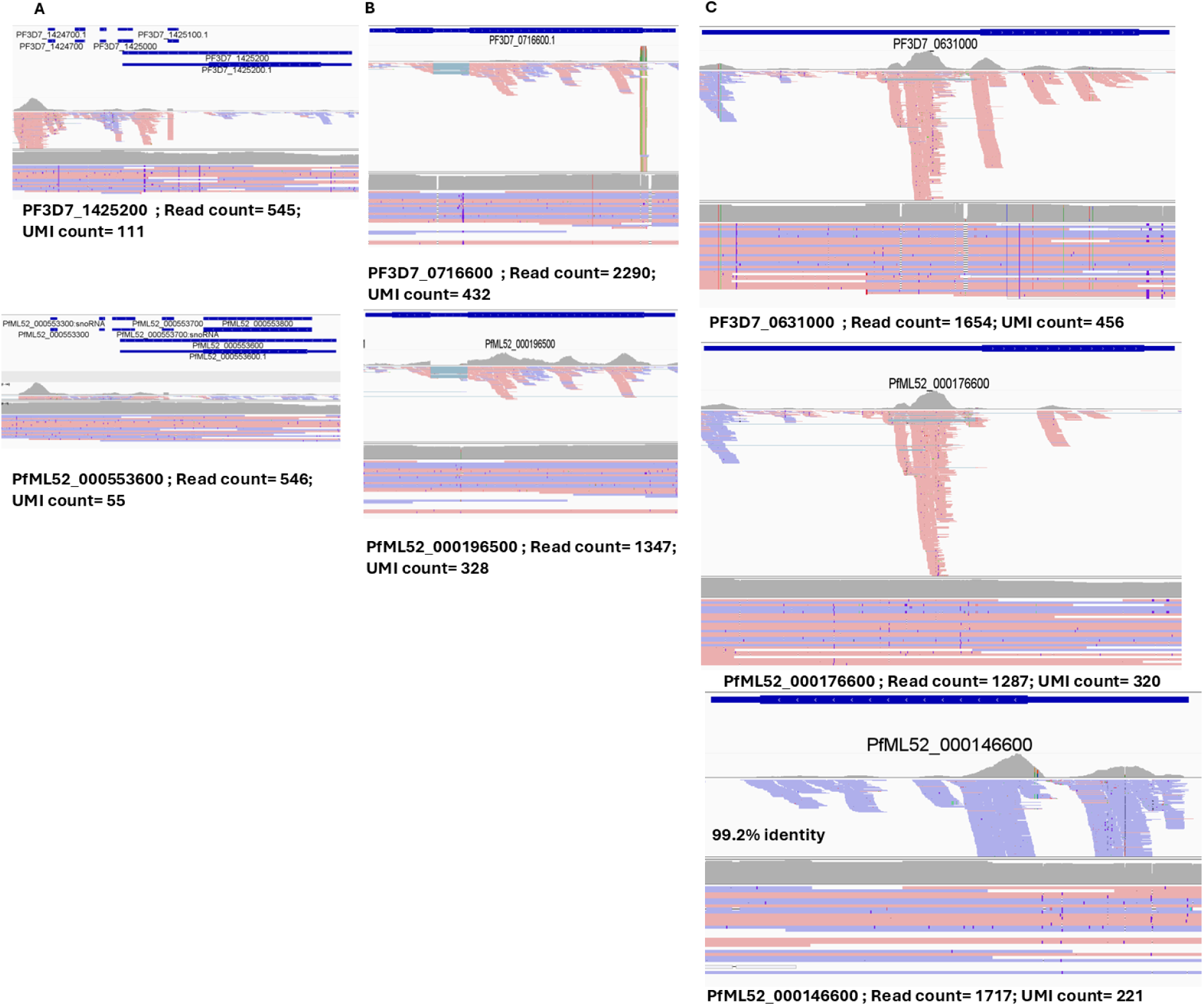
Locus-level sources of discordance in UMI-based quantification between PfML52 and Pf3D7 mappings. **(A)** Illustration of annotation-driven discrepancy at the *PfML52_000553600 / PF3D7_1425200* locus. In the PfML52 reference (bottom), this region contains two annotated isoforms/gene models (*PfML52_000553600* and *PfML52_000553800*), whereas the corresponding region in the Pf3D7 reference (top) lacks the gene without the UTRs (*PfML52_000553800*). As a result, reads are differentially allocated across features in PfML52 but aggregated in Pf3D7, leading to differences in UMI counts that are independent of true expression levels. **(B)** IGV visualization of the cysteine desulfurase locus (*PfML52_000196500 / PF3D7_0716600*) showing alignment irregularities in the Pf3D7 mapping (top), including fragmented coverage, mismatches, and positional shifts, compared to more coherent alignments in PfML52 (bottom). **(C)** IGV visualization of the tetratricopeptide repeat protein locus (*PfML52_000176600 / PF3D7_0631000*) showing reduced UMI counts in the PfML52 mapping relative to Pf3D7. A subset of reads assigned to the Pf3D7 locus was not recovered at the orthologous PfML52 locus but instead aligned to a closely related paralog (*PfML52_000146600*). However, this paralog was excluded from subsequent analyses because its reads had a median MAPQ of 3, indicating ambiguous multi-mapping, whereas reads assigned to the orthologous PfML52 gene were supported by uniquely mapped alignments (MAPQ = 255). Although the true origin of these reads cannot be determined with complete certainty because of the high sequence similarity between the loci, the mapping quality supports interpretation of the orthologous PfML52 locus as the more reliable assignment.

At the end of this analysis, five genes remained, comprising one SCO (*PfML52_000199500 / PF3D7_0719700*; 40S ribosomal protein S10, putative) and four non-SCO genes (*PfML52_000040100 / PF3D7_0215000*; and *PF3D7_0731600 / PfML52_000211500*; acetyl-CoA synthetase, *PfML52_000117700 / PF3D7_0508000*; 6-cysteine protein P38, and *PfML52_000298100 / PF3D7_1001100*; acyl-CoA binding protein ACBP1) that showed higher UMI counts in one reference relative to the other. For the remaining unresolved loci, several potential technical and biological sources of discrepancy were systematically evaluated. These included differences in strain composition across cells to test different parasites contributing reads differently, reference-dependent variation in gene assignment(GX/GN), and underlying genomic structure assessed using long-read alignments. In addition, Cell Ranger barcode and UMI tags (CB, UR, and UB) were examined to determine whether discrepancies originated from barcode assignment, or UMI correction and collapsing. None of these analyses provided a consistent explanation for the residual quantification differences.

## Multigene family analysis and reference-dependent mapping behaviour

### Gene set definition and family-level overview

Analysis of multigene families revealed a distinct pattern compared with the overall genome. In fact, in *P. falciparum*, multigene families such as *var*, *rifin*, *stevor*, *phist*, *etramp*, and *exported protein* families constitute a major component of the subtelomeric genome and encode proteins involved in antigenic variation and host–parasite interactions. We therefore performed a dedicated analysis of these families. Genes within each family were classified into three categories: (i) shared genes: Shared genes comprised all PfML52 genes for which Companion assigned a corresponding Pf3D7 gene identifier, irrespective of sequence identity, or mapping quality. No minimum sequence identity threshold was applied. For example, shared var genes exhibited sequence identities ranging from approximately 0.5 to 0.8. Consequently, highly polymorphic multigene family members, including var genes, were included despite frequently lacking strict one-to-one orthologous relationships. ; (ii) PfML52-specific genes lacking a corresponding Pf3D7 identifier; and (iii) Pf3D7-specific genes absent from the PfML52 annotation.

Because multigene families contain highly similar paralogous loci, reads classified as uniquely mapped (MAPQ = 255) may still originate from homologous genomic regions that cannot be fully resolved using short-read sequencing alone. For this reason, unlike the conserved core-gene analyses that were restricted to MAPQ = 255 reads, multigene family analyses incorporated both uniquely mapped and multi-mapped reads where appropriate. Restricting analyses exclusively to uniquely mapped reads would systematically exclude a biologically relevant fraction of reads originating from highly similar loci. Family-level quantification profiles were first evaluated using raw UMI counts visualized by scatter plots and heatmaps. Expression values were aggregated by developmental stage (early ring, late ring, early trophozoite, developing gametocyte, canonical female gametocyte, non-canonical female gametocyte, and male gametocyte), and profiles were compared between mappings to the ML52 genome and the Pf3D7 reference. Across most multigene families, expression profiles remained broadly concordant between references when the genes were considered to be found in both references (Fig. 10).

**Figure 10.**
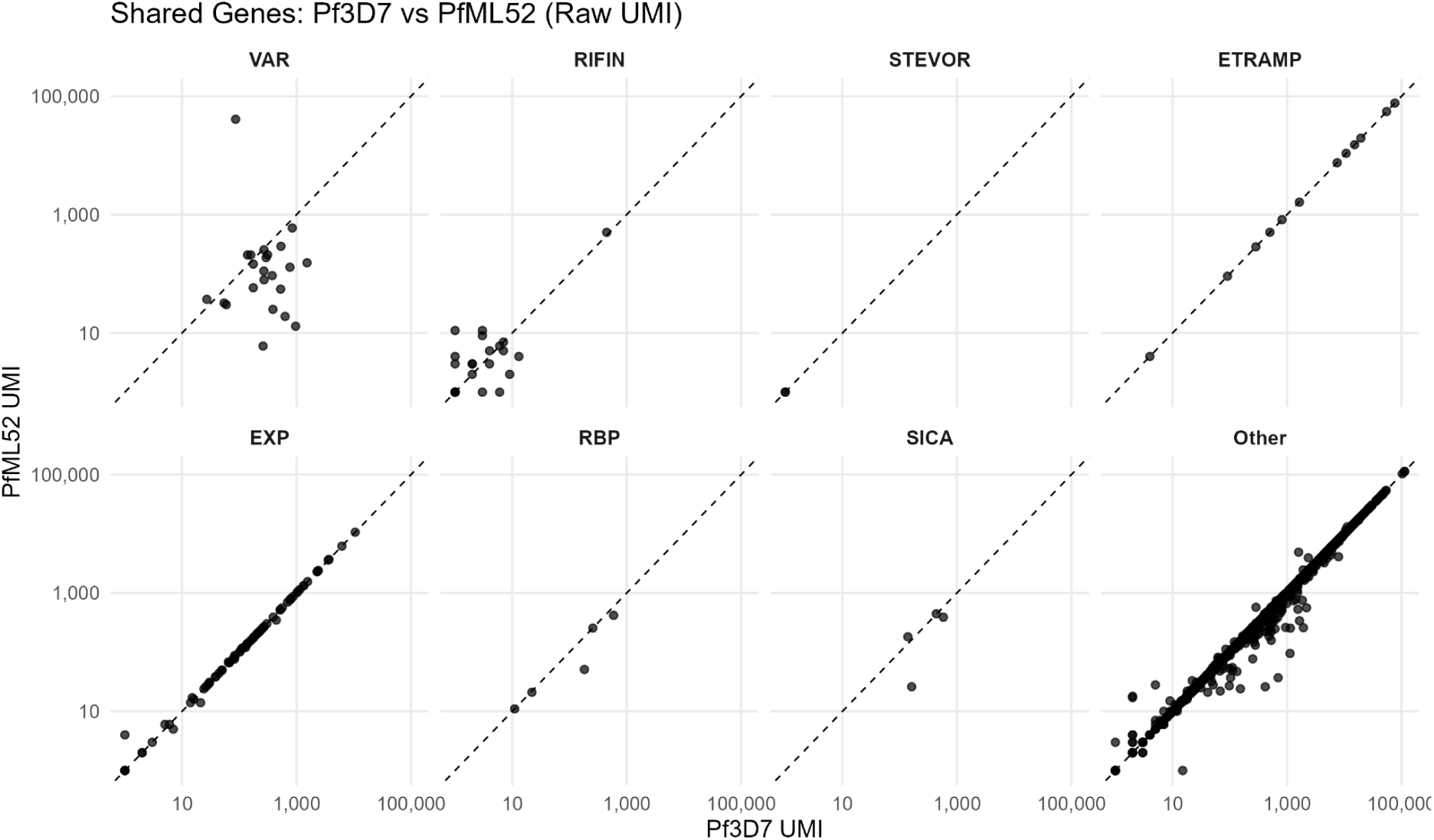
Scatterplots of raw UMI counts across shared genes of multigene families in PfML52 and Pf3D7. Encircled points in the *var* gene family indicate the *var* genes with the highest transcriptomic inference.

In contrast, pronounced reference-dependent differences were observed within the *var* gene family, where several highly expressed PfML52 *var* genes detected using the isolate-specific reference displayed markedly different expression profiles when the same reads were aligned to Pf3D7 (Fig. 11A, B). Overall, 406,324 reads aligned to *var* genes annotated as PfEMP1 in PfML52, compared with 138,793 reads aligned to Pf3D7 *var* genes. Among these, 128,351 reads were shared between references, whereas 277,973 reads aligned exclusively to PfML52 *var* loci and 10,442 aligned exclusively to Pf3D7 *var* loci (Fig. 11C). This latter category was unexpected and investigated further. Of the Pf3D7-exclusive reads, 2,190 mapped to unplaced ML52 contigs, while 2,078 exhibited MAPQ values below 255, indicating ambiguous or multi-mapping alignments. Together, these patterns demonstrate substantial reference-dependent read assignment within the *var* repertoire.

**Figure 11.**
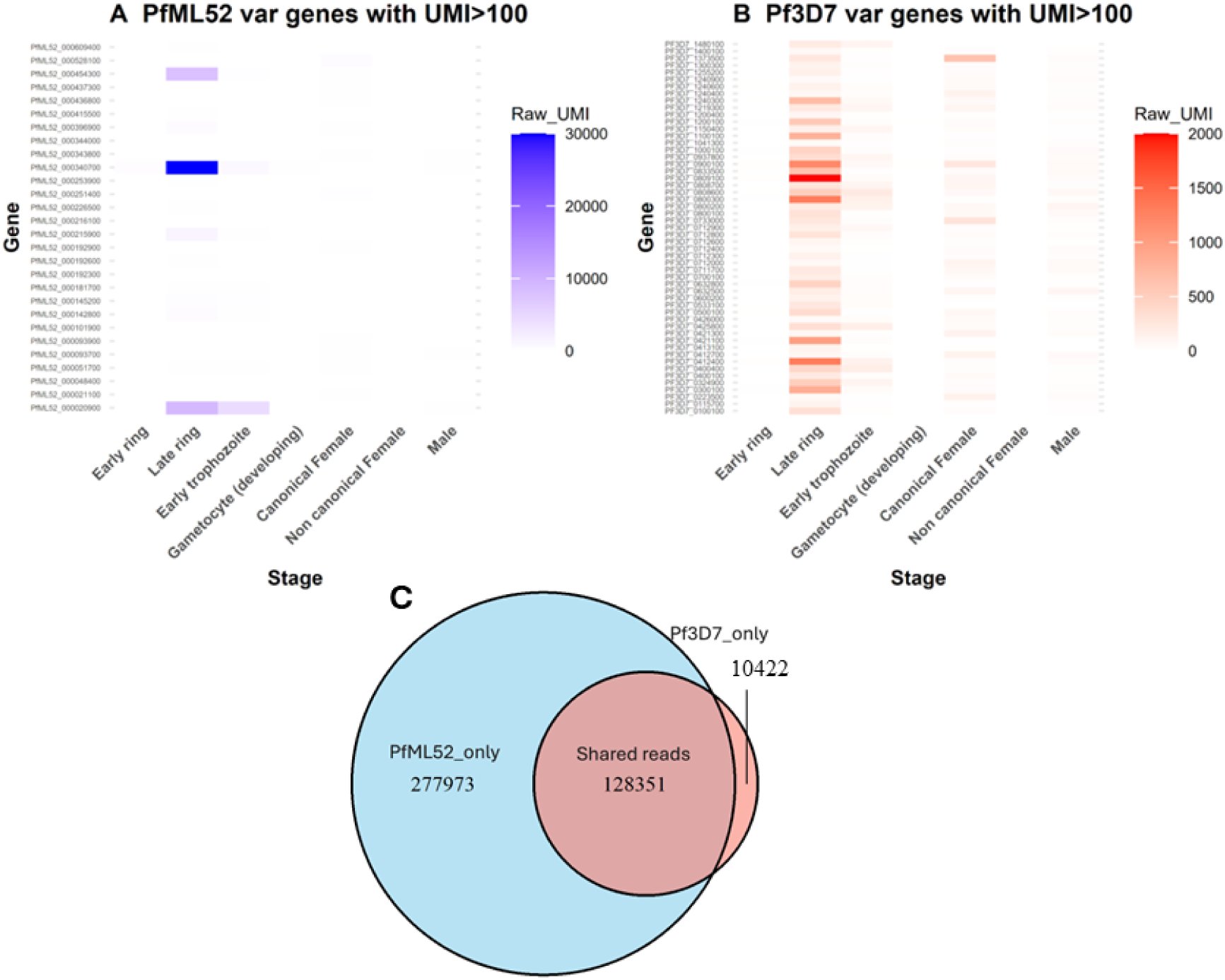
Reference-dependent distribution of var gene expression and read assignment between the PfML52 and Pf3D7 genomes. Heatmaps show all *var* genes in (A) PfML52 and (B) Pf3D7 with UMI counts >100. (C) Venn diagram showing the distribution of reads aligned to *var* loci in the PfML52 and Pf3D7 references. Most reads were assigned exclusively to PfML52 *var* loci, whereas reads assigned to Pf3D7 *var* loci were distributed across a larger number of genes.

We next examined the subset of reads observed in both references to determine whether they remained associated with corresponding loci or were redistributed among paralogous *var* genes.

To assist that, read-sharing relationships between PfML52 and Pf3D7 *var* genes were examined using bipartite gene–gene matrices, heatmaps, and alluvial visualisations. Analyses were restricted to reads that aligned to annotated *var* genes in both references, irrespective of whether they aligned to the same gene.

Analyses were performed at two levels. First, the complete read-sharing landscape was explored using all shared reads without filtering. Second, to focus on dominant relationships, only PfML52–Pf3D7 gene pairs connected by more than 500 shared reads were retained. This threshold was applied to individual pairwise relationships rather than to genes themselves; consequently, a single PfML52 *var* gene could remain connected to multiple Pf3D7 *var* genes if each relationship exceeded the threshold.

Analysis of *var*-gene mappings revealed two principal behaviours. First, many reads assigned to highly expressed PfML52 *var* genes were not recovered at annotated Pf3D7 *var* loci, indicating substantial reference-dependent read loss. Second, among reads aligned to *var* genes in both references, read assignments were frequently redistributed across multiple Pf3D7 loci rather than remaining associated with a single corresponding gene. This behaviour was particularly evident for *PfML52_000340700*, *PfML52_000020900*, and *PfML52_000454300*, which exhibited one-to-many relationships with multiple Pf3D7 *var* genes (Fig. 12A, B, Table. 3).

**Figure 12.**
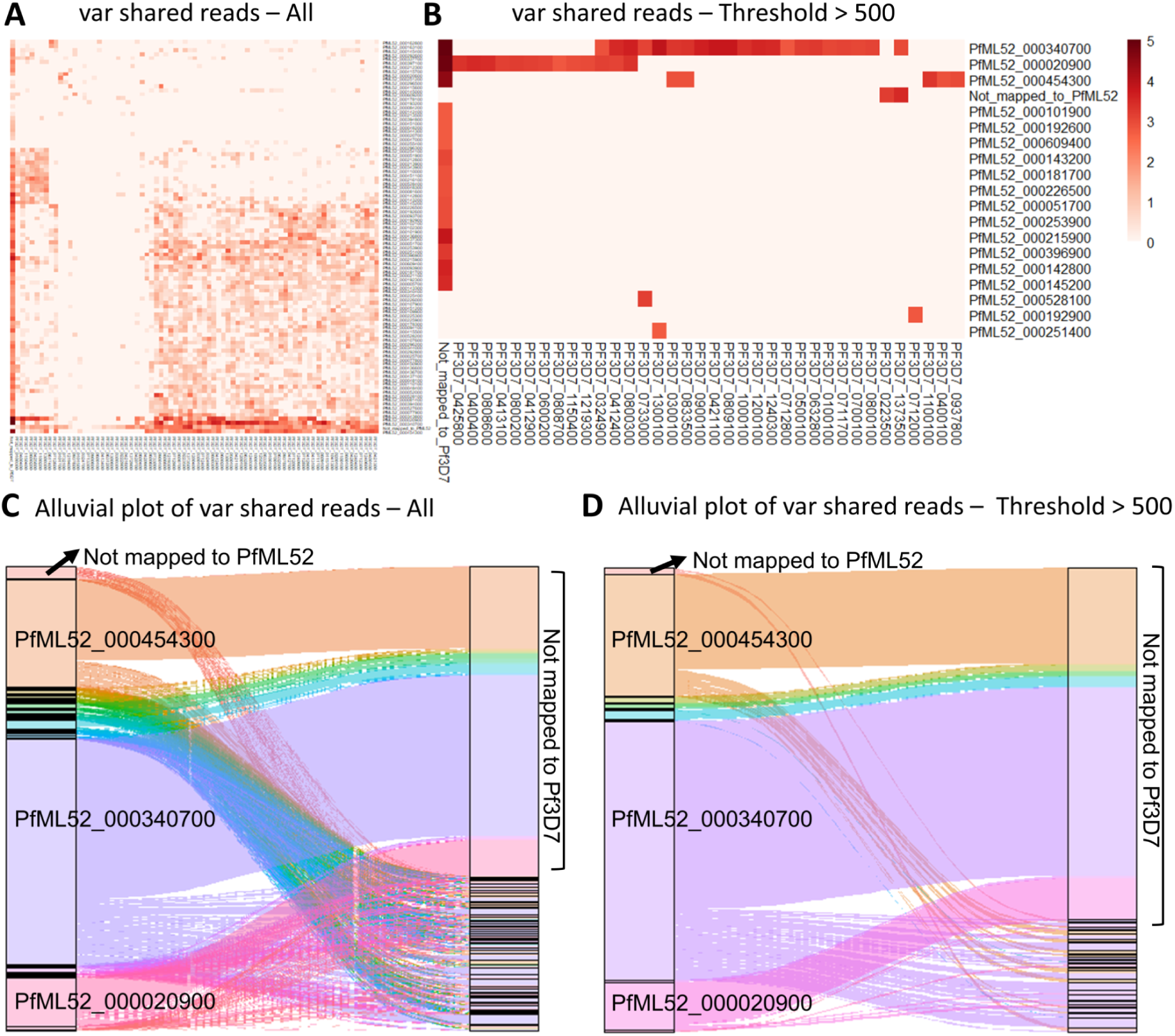
Read-sharing relationships between PfML52 and Pf3D7 *var* genes. **(A)** Heatmap showing pairwise relationships between PfML52 *var* genes (rows) and Pf3D7 *var* genes (columns) using all reads that are aligned to annotated *var* genes in both reference mappings. Here, shared reads refer to the same sequencing reads being aligned to *var* genes in both references and do not imply alignment to the same gene or to orthologous loci. Heatmap intensity represents log10(shared-read count + 1). **(B)** Heatmap showing dominant pairwise relationships after retaining only PfML52–Pf3D7 gene pairs connected by more than 500 shared reads. The threshold was applied to individual gene-pair relationships; consequently, a single PfML52 gene may be connected to multiple Pf3D7 genes. **(C)** Alluvial plots showing how reads from three highly expressed PfML52 *var* genes (*PfML52_000454300*, *PfML52_000340700*, and *PfML52_000020900*) were assigned across Pf3D7 *var* genes. Ribbon width is proportional to the number of shared reads connecting each PfML52–Pf3D7 gene pair. The complete set of pairwise relationships is shown without threshold filtering. **(D)** Alluvial plots showing dominant read-sharing relationships after retaining pairwise connections supported by more than 500 shared reads. “Not mapped to Pf3D7” indicates reads assigned to PfML52 *var* genes that lacked an annotated Pf3D7 *var*-gene alignment, whereas “Not mapped to PfML52” indicates reads assigned to Pf3D7 *var* genes that lacked an annotated PfML52 *var*-gene alignment.

This extensive one-to-many mapping pattern is consistent with the known architecture of *var* genes encoding *P. falciparum* erythrocyte membrane protein 1 (PfEMP1). PfEMP1 proteins contain conserved structural elements including an N-terminal segment (NTS), Duffy Binding-Like domains (DBL α–ε), cysteine-rich interdomain regions (CIDR α–γ), a transmembrane domain, and a conserved acidic terminal segment (ATS) [57]. Despite this conserved structural organization, *var* genes exhibit extensive sequence divergence, copy number variation, and recombination across parasite isolates. This diversity is thought to be maintained through frequent gene conversion and recombination events, generating highly diverse repertoires of PfEMP1 variants within natural parasite populations [58,59]. Consequently, short reads originating from conserved sequence blocks, particularly within exon 2 or conserved domain regions, can align to multiple paralogous loci in the reference genome.

Coverage inspection further supported expression from the three *var* loci in the PfML52 genome. Alignment of the scRNA-seq reads to the PfML52 genome produced continuous coverage across both exons of each gene, with exon–exon junction reads and no evidence of systematic exon-specific coverage bias. In contrast to the fragmented coverage observed when reads were aligned to the Pf3D7 reference, this continuous coverage supports transcription from these loci. Inspection of the splice junctions in IGV confirmed that all three genes used the expected canonical splice-site motifs, comprising GT–AG for the two forward-strand genes (*PfML52_000020900* and *PfML52_000454300*) and the complementary CT–AC motif for the reverse-strand gene (*PfML52_000340700*). The observed splice junctions coincided with the annotated intron boundaries, indicating that transcripts from all three loci were spliced at the annotated junctions.

Unexpectedly, antisense exon–exon junction reads were also observed for these three genes (Fig. 13, right). Antisense exon–exon junction reads represented 20.7% (25/121) and 6.1% (2/33) of all exon–exon junction reads for *PfML52_000020900* and *PfML52_000454300*, respectively, whereas they accounted for 55.0% (94/171) for the reverse-strand gene *PfML52_000340700*. To investigate whether these reads could result from Cell Ranger template-switching oligo (TSO) orientation mis-assignment, we compared template-switch trimming (ts:i) values between sense-spliced, sense-unspliced, antisense-spliced and antisense-unspliced reads for each gene. Across all three genes, antisense-classified spliced reads showed a similar distribution of ts:i values to sense-spliced reads and used the same genomic splice junctions as the sense transcripts, which are canonical only in the sense orientation. These observations suggest that the antisense-classified spliced reads may reflect cDNA-derived orientation artefacts rather than genuine antisense splicing. Consistent with this interpretation, antisense reads were distributed throughout the gene bodies, as illustrated by the mixed blue and red reads in each IGV visualisation, rather than forming distinct antisense transcript structures.

**Figure 13.**
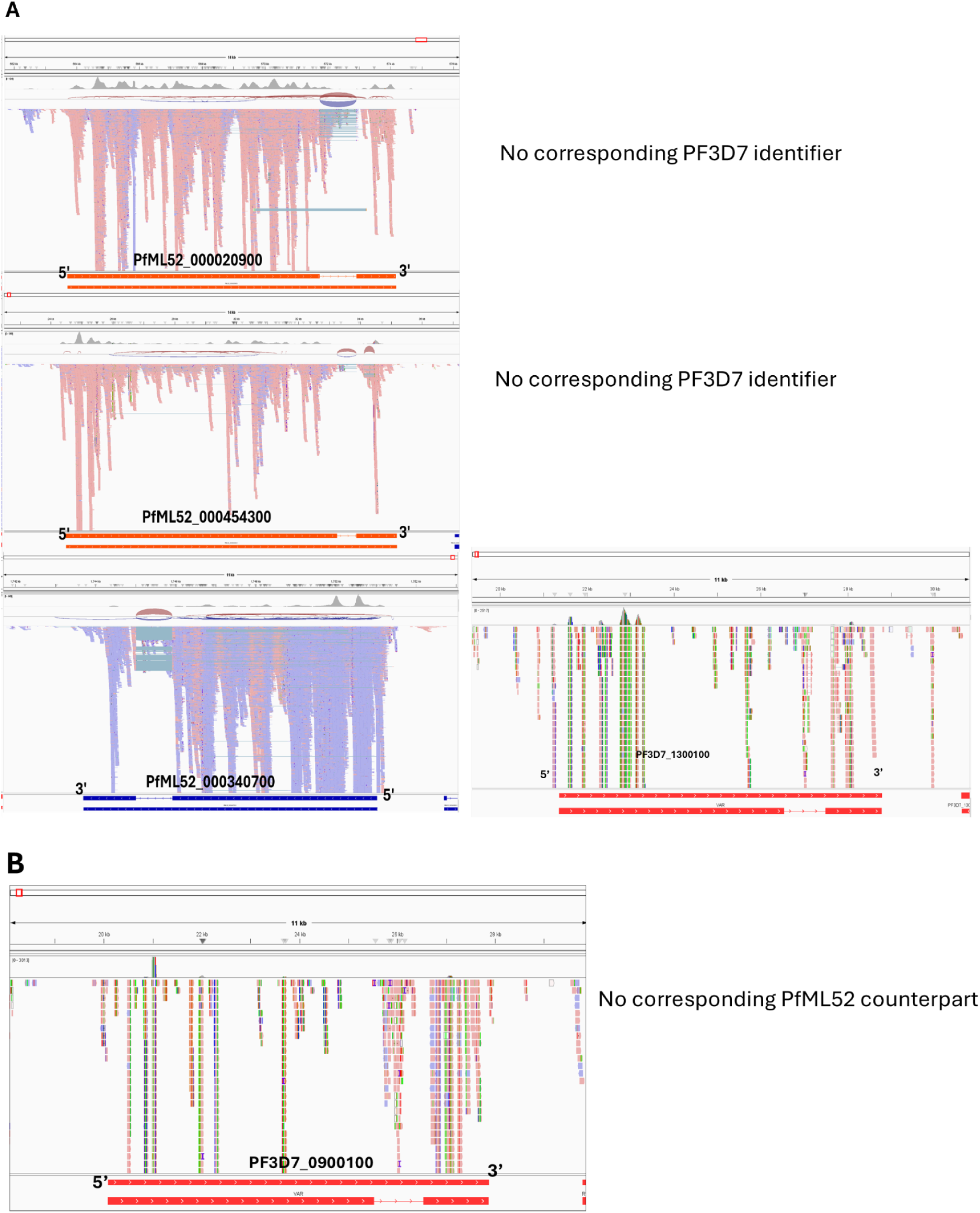
IGV inspection of short-read alignments across the highest expressed var loci in the PfML52 and Pf3D7 reference genomes. IGV inspection of short-read alignments across the top three *var* genes with the highest UMI counts identified in the PfML52 reference and their Companion-assigned Pf3D7 orthologues, where present (A), and the top *var* gene in Pf3D7 *PF3D7_0900100* in the Pf3D7 reference (B). Reads aligned to the isolate-specific PfML52 reference show continuous coverage across individual var loci, whereas alignment to the Pf3D7 reference redistributes reads across multiple var genes, resulting in fragmented coverage at individual loci. Gene models are shown beneath each locus. Reads are coloured according to their alignment strand (red and blue), and gene annotations indicate genes encoded on the forward or reverse genomic strand. Of the three PfML52 var genes, only *PfML52_000340700* has an orthologue in the Pf3D7 reference (*PF3D7_1300100*); *PfML52_000020900* and *PfML52_000454300* have no annotated orthologues in Pf3D7.

Together, these findings indicate that the three dominant *var* loci produce correctly spliced transcripts. In contrast, mapping to Pf3D7 produced fragmented coverage profiles, and reduced read support. (Fig. 13, B). These findings indicate that apparent reference-dependent expression differences primarily reflect mistaken mapping of reads to non-orthologous/homologous *var* loci rather than genuine differences in expression inference. More broadly, the results suggest that isolate-specific genome assembly can recover biologically coherent *var* expression patterns that are obscured, or worse – incorrect, when reads are mapped to a divergent reference repertoire. They also highlight the value of pseudobulk analyses for *var* gene investigations, where aggregating reads across cells can increase support across gene bodies and facilitate interpretation of highly polymorphic multigene families.

A subset of reads appeared to align uniquely to Pf3D7 *var* genes without a retained alignment to the PfML52 assembly. Manual inspection showed that these were not independent Pf3D7-specific transcripts, but represented subsets of larger groups of reads distributed across homologous *var* loci. For example, PF3D7_1373500 was supported by 5,153 aligned reads, of which 2,943 lacked a retained alignment to the PfML52 assembly after alignment filtering, whereas the remaining 2,210 also aligned to homologous PfML52 *var* genes. Similarly, PF3D7_0100100 was supported by 1,823 reads, of which only 82 lacked a retained PfML52 alignment, while the remainder aligned across 52 homologous PfML52 *var* loci. Comparable patterns were observed for 72 Pf3D7 *var* genes.

These observations are consistent with reference-dependent read assignment in highly repetitive var regions. One possible explanation is that homologous regions are represented differently in the two reference genomes, with some PfML52 var loci being fragmented, partially unresolved, or located on unplaced contigs. Under these circumstances, reads originating from conserved var regions may preferentially retain alignments to Pf3D7 loci when equivalent PfML52 alignments are not retained by the aligner. Importantly, MAPQ = 255 in this context does not necessarily indicate biological uniqueness, but rather the absence of retained alternative alignments within the supplied reference. Consistent with this interpretation, many apparently Pf3D7-exclusive reads either mapped to unplaced PfML52 contigs or exhibited lower mapping quality scores characteristic of ambiguous alignments. Because these analyses were performed on reads from all parasite populations, we also cannot exclude a contribution from parasites belonging to strains other than SC2. Together, these observations suggest that much of the apparent Pf3D7-specific *var* expression is likely to reflect reference-dependent alignment behaviour rather than genuine strain-specific transcription.

### Stage-specific expression of the PfML52 *var* repertoire

To investigate whether *var* gene expression was associated with the parasite developmental stage, we examined all PfML52 *var* genes with total expression exceeding 100 UMIs. For each gene, we calculated the proportion of total transcript abundance originating from asexual parasites (early ring, late ring and early trophozoite) and sexual parasites (female and male gametocytes).

The analysis revealed marked heterogeneity among individual *var* genes. The most highly expressed *var* genes were almost exclusively expressed during asexual development, with less than 5% of their total transcript abundance detected in sexual stages. In contrast, several moderately expressed *var* genes exhibited were predominantly expressed during sexual stages, while a small number of genes appeared to derive most of their transcript abundance from gametocytes.

To determine whether these apparently sexually expressed var genes represented genuine transcription rather than mapping artefacts, all candidate loci were manually inspected using the Integrative Genomics Viewer (IGV).

One representative example was *PfML52_000528100*, which initially appeared to be highly enriched in female gametocytes, with 497 of 571 total UMIs (87.0%) originating from female-stage cells. However, manual inspection in IGV showed that the mapped reads were restricted to short conserved regions rather than spanning the gene continuously. Although these reads carried MAPQ = 255 and were therefore assigned uniquely by the aligner, they likely originate from highly conserved sequence shared among closely related *var* genes.

Consequently, the observed female-stage signal cannot be considered definitive evidence of genuine transcription of the complete *PfML52_000528100* locus.

Overall, our data support the conclusion that the major transcriptional output of the PfML52 *var* repertoire remains associated with asexual parasites.

### Mapping to the isolate-specific PfML52 genome reveals strain- and stage-associated *var* expression patterns

To investigate *var* gene expression using the isolate-specific PfML52 reference genome, we examined single-cell expression of the three most highly expressed *var* genes identified in the dataset (*PfML52_000340700*, *PfML52_000020900* and *PfML52_000454300*) across all parasite strains and developmental stages.

A heatmap of single-cell expression demonstrated that the three dominant PfML52 *var* genes were expressed predominantly within the SC2 parasite population and were largely associated with asexual developmental stages, including early rings, late rings and early trophozoites (Fig. 14). In contrast, these genes showed only low expression in SC3 and SC4 parasites and were largely absent from SC1 and SC5. Little or no expression of these three PfML52 *var* genes was detected in the sexual-stage parasites. However, because the sexual parasites belong to a different genetic background, this observation should not be interpreted as evidence for globally reduced *var* expression during sexual development, but rather indicates that these specific ML52 *var* genes are not prominently represented in that population. Together, these observations indicate that expression of the dominant PfML52 *var* genes is largely restricted to the SC2 asexual parasite population, supporting the subsequent detailed analysis of *var* transcription in this strain.

**Figure 14.**
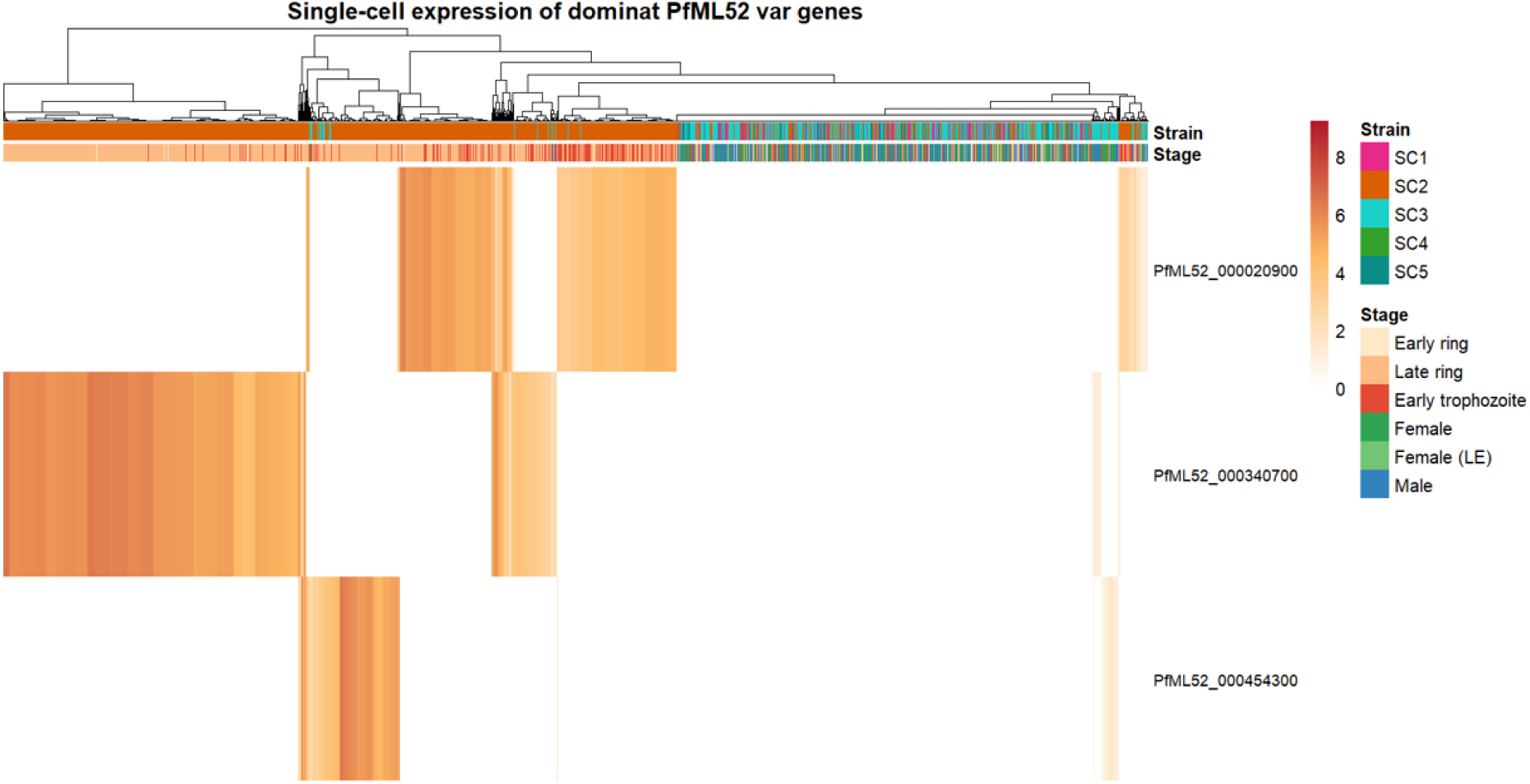
Heatmap of single-cell expression of the three dominant PfML52 *var* genes across parasite strains and developmental stages. Heatmap showing log-normalized single-cell expression of *PfML52_000020900*, *PfML52_000340700* and *PfML52_000454300* across five *P. falciparum* strains and both asexual and sexual developmental stages. Columns represent individual cells annotated by strain and developmental stage, and rows correspond to the three dominant *var* genes. Colour intensity indicates normalized expression level, with darker colours representing higher expression. Expression of all three genes is predominantly restricted to the SC2 asexual parasite population.

To provide developmental context for SC2 parasites, we classified the asexual-stage parasites into early-ring, late-ring and early-trophozoite stages, which represent a continuous developmental trajectory. This framework was used to assess stage-specific *var* expression patterns.

### Single-cell analysis identifies three major *var* expression programmes in the SC2 parasite population

We next examined expression of the three dominant *var* genes within SC2 parasites (n = 6,567 cells). *PfML52_000340700* was detected in 3,060 cells (46.6%), *PfML52_000020900* in 2,153 cells (32.8%), and *PfML52_000454300* in 793 cells (12.1%). Most parasites expressed only a single dominant *var* gene. Specifically, 2,803 cells expressed only *PfML52_000340700*, 1,912 expressed only *PfML52_000020900*, and 670 expressed only *PfML52_000454300*. A further 309 parasites expressed two dominant *var* genes, comprising 187 cells expressing *PfML52_000020900* and PfML52_000340700, 69 cells expressing *PfML52_000340700* and *PfML52_000454300*, and 53 cells expressing *PfML52_000020900* and *PfML52_000454300*. Only a single parasite expressed all three dominant *var* genes, whereas 872 cells showed no detectable expression of these three loci.

To identify the predominant *var* transcript in each parasite, every cell was assigned to the *var* gene with the highest normalized expression level among the three dominant PfML52 *var* genes. This analysis identified three major *var*-expression programmes within SC2, comprising 2,947 parasites dominated by *PfML52_000340700* (44.9%), 2,024 dominated by *PfML52_000020900* (30.8%), and 724 dominated by *PfML52_000454300* (11.0%). This approach identifies the dominant transcript in each cell and does not imply exclusive expression of a single *var* gene; parasites expressing multiple dominant *var* genes were analysed separately below. Projection of these expression classes onto the UMAP revealed partially overlapping distributions across the developmental trajectory, indicating that the three dominant *var* programmes represent distinct transcriptional states that may be associated with parasite development rather than discrete cell clusters. (Fig. 15).

**Figure 15.**
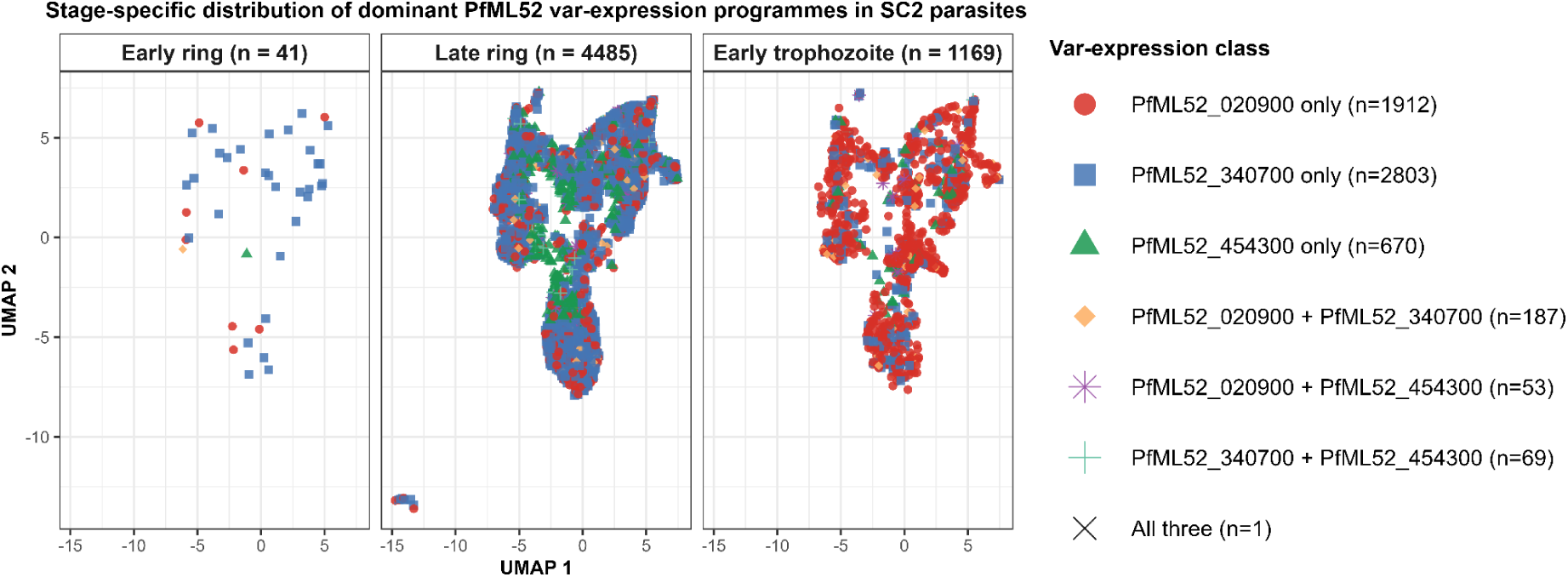
Stage-specific distribution of dominant PfML52 *var*-expression programmes in SC2 parasites.

**Figure 16.**
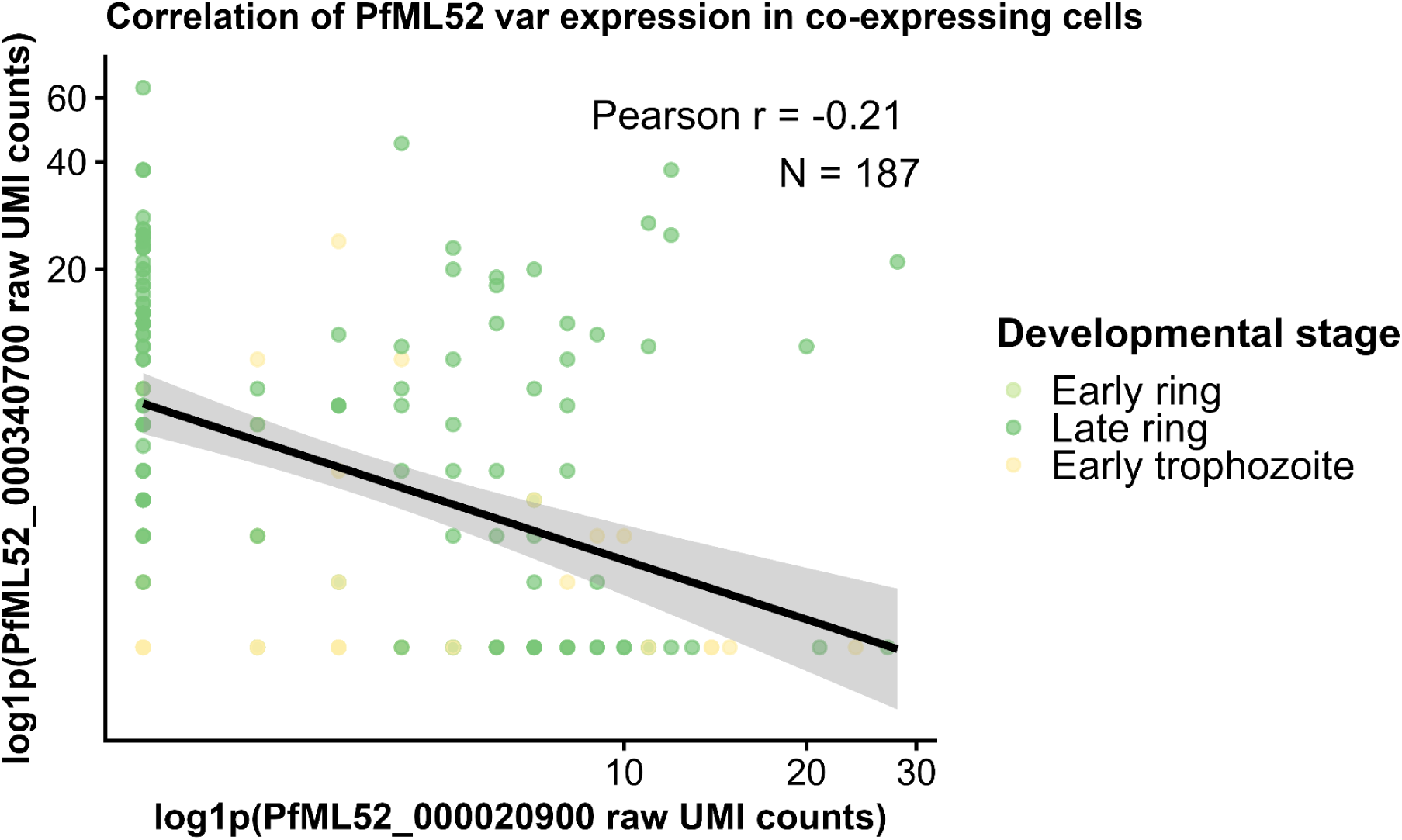
Correlation of PfML52_000020900 and PfML52_000340700 expression in SC2 cells co-expressing both var genes. Scatter plot showing the relationship between the expression of PfML52_000020900 and PfML52_000340700 in SC2 cells co-expressing both var genes. Each point represents one cell. N: number of co-expressing cells.

Together, these findings demonstrate that mapping reads to the matched isolate-specific PfML52 reference genome resolves three biologically distinct and developmentally associated *var* expression programmes within the SC2 parasite population (Fig. 14). Although most parasites expressed a single dominant *var* gene, consistent with the principle of mutually exclusive *var* transcription, the dominant gene changed with parasite development: *PfML52_000340700* and *PfML52_000454300* predominated in late-ring parasites, whereas *PfML52_000020900* extended into early trophozoites. The small subset of dual-positive parasites was largely restricted to later developmental stages and may represent transitional states during *var* switching rather than stable co-expression of multiple dominant *var* genes. These patterns were readily detectable when reads were aligned to the isolate-specific reference genome but were obscured when mapping was performed against the divergent Pf3D7 reference genome.

Overall, our findings demonstrate that the impact of reference genome choice depends strongly on genomic context. Expression inference for conserved core genes remained largely stable between references, whereas *var* genes showed substantial sensitivity. Integrating isolate-specific genome assemblies with read-level analyses and locus-level validation provides an effective framework for distinguishing technical mapping artefacts from genuine biological signals. Such genome-aware approaches are likely to be particularly important for pathogens characterized by extensive antigenic variation systems and highly dynamic subtelomeric genomes, including *P. falciparum*, *Trypanosoma brucei* [60], *Borrelia burgdorferi* [61], and *Neisseria gonorrhoeae* [62].

However, some limitations should be considered. This study analysed a single patient isolate and was designed as a mechanistic investigation rather than a population-scale survey. Although the specific loci affected by mapping artefacts may differ between isolates, the underlying drivers—including sequence divergence, copy number variation, paralogous gene content, and annotation ambiguity—are intrinsic features of *P. falciparum* circulating strains and are therefore likely to be relevant across genetically diverse infections.

An additional consideration is the multiclonal nature of the ML52 infection, which contained five parasite haplotypes. The presence of multiple haplotypes increases the difficulty of genome assembly and read mapping, particularly within repetitive and subtelomeric regions. Divergent haplotypes may assemble into separate contigs or remain partially unresolved, potentially generating apparent mapping asymmetries when reads derived from minor variants align suboptimally to the dominant haplotype represented in the assembly. Some unplaced contigs may therefore reflect divergent haplotypes and/or unresolved repetitive subtelomeric sequence. Nevertheless, multiclonal infections are characteristic of *P. falciparum* transmission in endemic settings, making this dataset representative of biologically realistic field conditions rather than an artificial experimental system.

Despite these limitations, the overall conclusions remain consistent. Short-read scRNAseq expression inference in *P. falciparum* is largely robust to reference genome variation for conserved and single-copy loci. In contrast the *var* repertoire shows substantial sensitivity to reference genome and alignment context, leading primarily to redistribution of reads among partial homologous loci rather than true expression inference. Integrating isolate-specific genome assembly with genome-aware mapping and locus-level validation therefore provides a robust framework for interpreting transcriptomic data from genetically diverse malaria parasites.

## Supporting information

Table S1. Annotation table and annotation sources for the PfML52 genome relative to the Pf3D7 genome.

Table S2. Gene_ids of twelve genes in PfML52 and their orthologs in Pf3D7 with multiple annotation issues.

## Reproducibility and data availability

All mapping, assembly, and annotation pipelines are publicly available (see Data Availability section). Analysis scripts and processed data tables for gene-level metrics and mapping quality are included in the Supplementary Materials.

## Abbreviations

scRNA-seq: single-cell RNA sequencing
UMI: unique molecular identifier
MAPQ: mapping quality
PfML52: a patient-derived *P. falciparum* isolate obtained from a Malian patient
CPP: Conserved Plasmodium Protein
PHIST-RESA: Plasmodium Exported Protein Ring-Infected Erythrocyte Surface Antigen
ETRAMP: Early Transcribed Membrane Protein
RIFIN: Repeat Interspersed Family
STEVOR: Subtelomeric Variable Open Reading Frame
SICA: C-terminal Inner Membrane domain containing protein
HYPO: hypothetical proteins
Rest: other genes.
PfEMP1: *P. falciparum* erythrocyte membrane protein 1

## Funding

All authors and the work contained within were funded by Wellcome award 220540/Z/20/A to the Wellcome Sanger Institute. Additionally, the scRNAseq data was generated as part of an MRC funded project to MKNL (MR/S02445X/1).

## Acknowledgments

The author acknowledges the support of Cara (the Council for At-Risk Academics), which provided fellowship support to the author during this work. We would like to thank Petra Korlevic and Alex Makunin for their valuable comments, critical review of the manuscript, and constructive suggestions that helped improve this work. The sample in this study from a natural infection was generated as part of a Medical Research Council grant MR/S02445X/1 to MKNL.

## Author contributions

Conceptualization: Sunil Kumar Dogga & Mara Lawniczak, methodology: Sunil Kumar Dogga & Mara Lawniczak, formal analysis: Talleh Almelli, writing—original draft: Talleh Almelli, writing—review & editing: Talleh Almelli, Sunil Kumar Dogga, Jesse Rop, Seri Kitada & Mara Lawniczak, supervision: Sunil Kumar Dogga & Mara Lawniczak

## Disclosure of use of AI-assisted tools including generative AI

ChatGPT (OpenAI, GPT-5.5) and Grammarly were used exclusively for language editing, grammar correction, and rephrasing of selected text passages. All scientific analyses, interpretations, figure generation, and conclusions were developed, verified, and approved by the authors.

## Data availability & Reproducibility

The genome assembly, genome annotation, processed single-cell expression matrices, and analysis scripts supporting this study are publicly available on Zenodo at https://doi.org/10.5281/zenodo.21622849. Raw single-cell RNA sequencing data have been deposited in the European Nucleotide Archive (ENA) and will become publicly available upon manuscript publication. The ENA accession number will be provided after release.

## Competing interests

The authors declare no competing interests.

---

**Figure S1:**
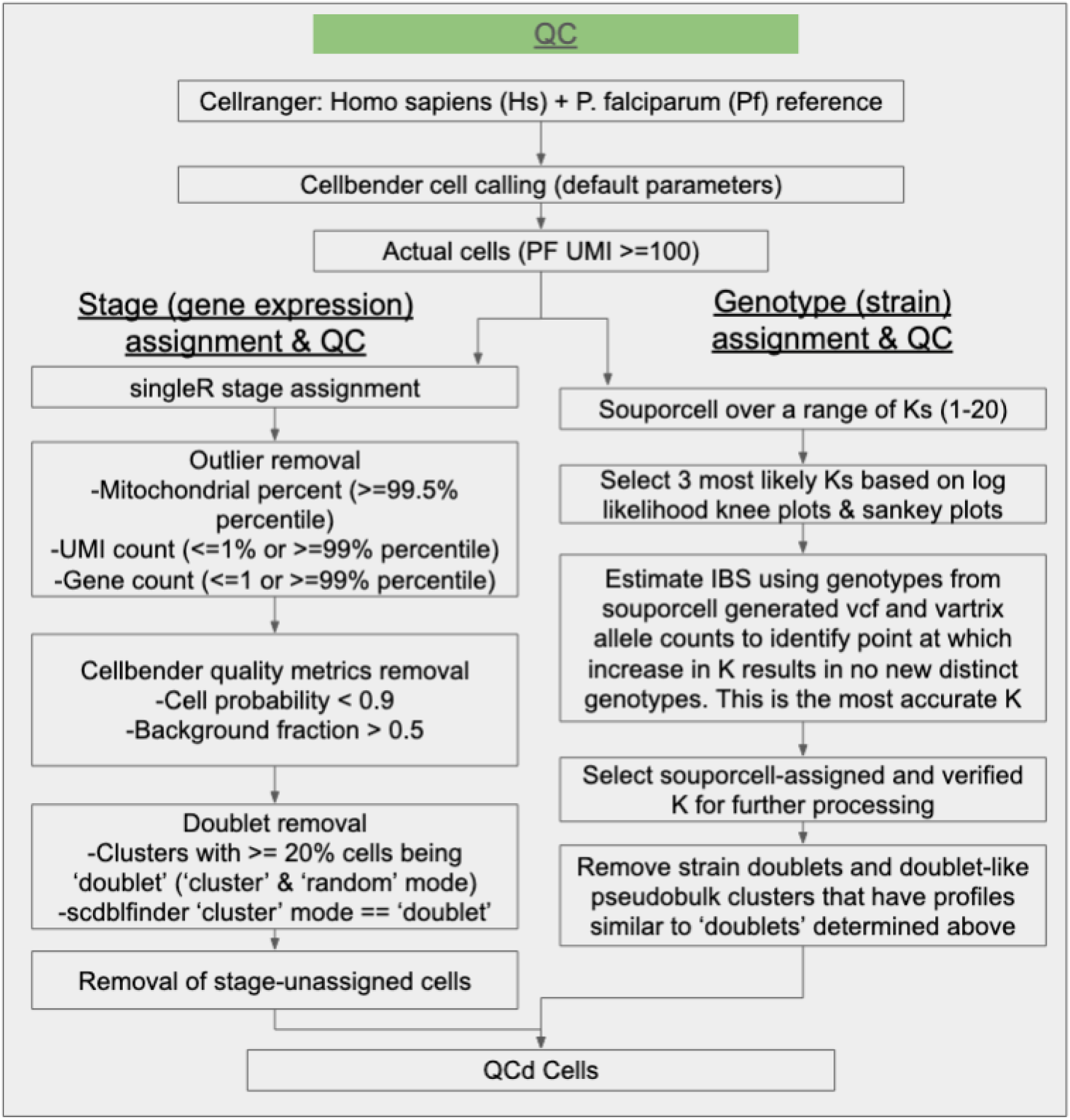
Workflow of the PfML52 natural infection QC process.

**Table S1.**
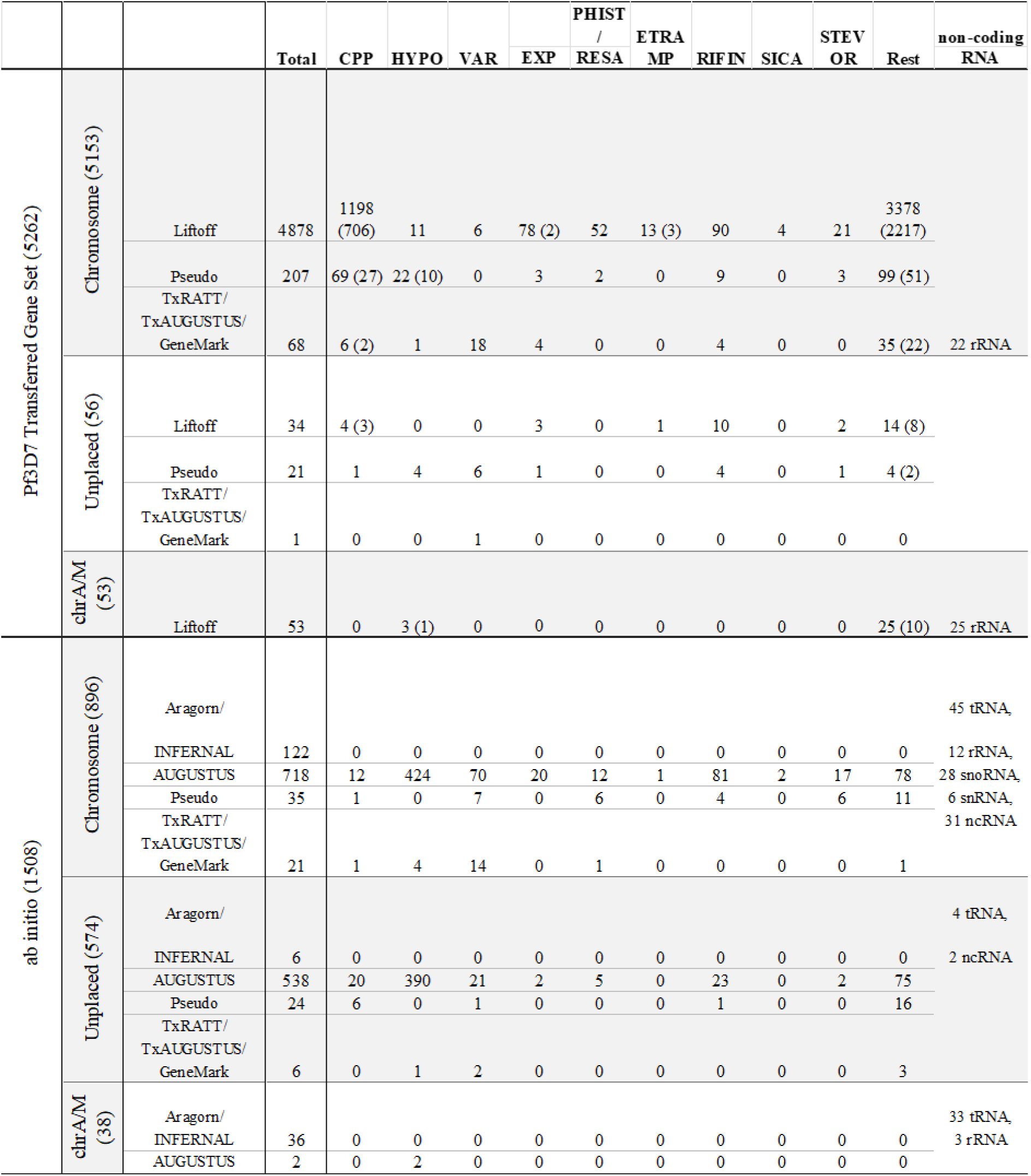
Annotation table and annotation sources for the PfML52 genome relative to the Pf3D7 genome. The table is divided into two major sections: (i) the *Pf3D7 transferred gene set*, comprising loci with identifiable Pf3D7 orthologs and reference-guided annotation support, and (ii) *ab initio annotations*, comprising loci predicted primarily through computational gene prediction approaches. Numbers in brackets indicate the number of single-copy ortholog (SCO) genes within each category. Within the transferred gene set, genes annotated by Liftoff represent high-confidence ortholog transfers in which Pf3D7 gene models were directly mapped onto the PfML52 assembly while preserving exon–intron structure and genomic synteny. “Pseudo” denotes putative pseudogenes identified during transfer, including loci with disrupted coding structure, frameshifts, premature stop codons, or fragmented alignments relative to Pf3D7. TxRATT/TxAUGUSTUS/GeneMark represent reference-guided reconstructed annotations in which direct transfer was incomplete or unsuccessful due to structural variation, sequence divergence, or assembly differences; in these cases, Companion used homology-assisted prediction methods to reconstruct gene models while retaining association with corresponding Pf3D7 orthologs and identifiers. Consequently, these loci may still have sequence identity and coverage values relative to Pf3D7 orthologs despite not being directly transferred by Liftoff. Within the *ab initio* section, AUGUSTUS and GeneMark correspond to computationally predicted coding genes generated primarily through homology-assisted gene prediction. These loci may include hypothetical genes lacking clear Pf3D7 orthologs as well as predictions supported by Pf3D7 protein evidence. Aragorn/Infernal were used for annotation of non-coding RNA genes, including tRNA, rRNA, snoRNA, snRNA, and ncRNA loci. CPP: Conserved *Plasmodium* Protein, HYPO: hypothetical proteins, PHIST-RESA: *Plasmodium* Exported Protein Ring-Infected Erythrocyte Surface Antigen, ETRAMP: Early Transcribed Membrane Protein, RIFIN: Repeat Interspersed Family, STEVOR: Subtelomeric Variable Open Reading Frame, SICA: C-terminal Inner Membrane domain containing protein, Rest: other genes.

**Table S2.** Gene_ids of twelve genes in PfML52 and their orthologs in Pf3D7 with multiple annotation issues.

| PfML52 gene_id | Pf3D7 gene_id |
| --- | --- |
| *PfML52_000013000* | *PF3D7_0108600* |
| *PfML52_000038200* | *PF3D7_0213400* |
| *PfML52_000283000* | *PF3D7_0928500* |
| *PfML52_000432900* | *PF3D7_1236900* |
| *PfML52_000482600* | *PF3D7_1327700* |
| *PfML52_000487600* | *PF3D7_1332600* |
| *PfML52_000553600* | *PF3D7_1425200* |
| *PfML52_000569000* | *PF3D7_1440200* |
| *PfML52_000012900* | *PF3D7_0108500* |
| *PfML52_000038100* | *PF3D7_0213200* |
| *PfML52_000155500* | *PF3D7_0610400* |
| *PfML52_000203900* | *PF3D7_0724100* |

## Notes

### Competing Interest Statement

The authors have declared no competing interest.

